# Global cell-state and gene-program representations reveal conserved and context-specific perturbation responses of cells

**DOI:** 10.64898/2026.05.16.725005

**Authors:** Xingjie Pan, Reuben Saunders, Joseph M. Replogle, Jonathan Weissman, Xiaowei Zhuang

## Abstract

Understanding how cell states change in response to genetic perturbations is critical for gene-function and therapeutics discovery. However, state-of-the-art deep-learning models trained on large single-cell omics datasets still struggle to accurately predict cellular responses to perturbations, highlighting the need for a better understanding of the cell-state space and how cells move through this space. Here, we present a contrastive learning model that integrates diverse scRNA-seq datasets into a global, interpretable cell-state manifold. We further develop a framework to integrate this global cell-state manifold with genome-scale perturbation data to identify gene-expression programs that define principal axes of cell-state transitions and functional embeddings of genes that define major perturbation classes. Applying this framework across Perturb-seq datasets on different cell types reveals conserved cellular responses to perturbations, as well as cell-type-specific rewiring of stress responses. Moreover, we perform a genome-scale Perturb-seq screen in human embryonic stem cells, validating and extending these findings and uncovering a class of mesenchymal transitions induced by diverse perturbations to cellular stress-response pathways.

## Introduction

Mammalian cells exhibit a broad spectrum of states, encompassing lineage-specified cell-type identities established during development, transient functional states, and pathological states arising in aging and disease. Advances in single-cell transcriptomics now make it possible to systematically profile these cell states at massive scale^1–6^, raising the prospect of constructing generalizable models that predict how cells respond to perturbations^7^. Because most functional cell biology experiments involve measurements of cellular responses to some form of perturbations (genetic, pharmacological, or environmental), such models could greatly accelerate biological discovery by enabling rapid in silico experimentation^2,7^. Inspired by progress in large language models, recent efforts have trained foundation models on large compendia of single-cell RNA-sequencing (scRNA-seq) datasets to learn gene- and cell-level embeddings that capture latent gene regulatory structures^8^. Notwithstanding the success in transfer-learning tasks, these models provide limited improvement in predicting gene-expression responses to genetic perturbations; notably, recent benchmarking studies indicate that complex deep-learning models did not outperform simple linear baselines in prediction accuracy^9^. This persistent gap between data abundance and prediction accuracy points to a major challenge in applying generic black-box approaches to perturbation modeling. Achieving progress—whether by developing better models or collecting more informative training datasets—thus requires a deeper understanding of the relationships between different cell states and the principles governing how cell states change in response to perturbations. This entails solving a number of key questions: What is the overall structure of the cell-state space? How do cells traverse this space in response to perturbations? What are the principal axes that organize cell-state transitions? Which classes of perturbations drive movement along each axis? And how does context, such as intrinsic cell state or extrinsic conditions, rewire cellular response to the same perturbation?

Addressing these questions would benefit from a global framework that moves beyond identifying isolated insights and places diverse single-cell genomics (e.g. scRNA-seq) and functional genomics (e.g. Perturb-seq) data into a common, interpretable coordinate system for comparative analyses of cell states, perturbations, and cellular responses to perturbations. Such framework should include: 1) a representation of the global cell-state landscape that captures its overall structure and allows new cell states to be mapped consistently, enabling direct comparison of cell states across datasets; 2) a way to reduce the high dimensionality of cell-state transitions into a set of principal axes such that the variations in hundreds to thousands of genes are reduced into a manageable and interpretable number of principal components; 3) a method for cross-context dimensionality reduction and classification of perturbations, such that the vast space of possible perturbations can also be reduced to principal axes that group perturbations eliciting similar cellular responses, which are shared across different intrinsic and extrinsic cellular context. Establishing such a framework would not only reveal the global structure of the spaces of cell states and perturbations, but also provide a foundation for generalizable analysis of diverse datasets and systematic understanding of cell-state transitions and perturbation responses.

Despite these advantages, constructing a framework that satisfies these goals is technically challenging. A central challenge in constructing a global and interpretable representation of cell states—and in performing consistent dimensionality reduction for cell-state transition—is the pervasive presence of batch effects in single-cell data. Advances in scRNA-seq have yielded highly diverse datasets that together span nearly the entire landscape of natural human and mouse cell states^2^, and existing dimensionality reduction methods can generate interpretable low-dimensional representations for individual datasets^10^. However, no single dataset captures the full diversity of cell states, and batch effects often separate biologically similar states from different experiments. Current integration methods can mitigate these effects for pairs of datasets with overlapping cell-state compositions by aligning their major axes of variation^11^, but they do not scale effectively to large, partially overlapping collections. Recent deep-learning approaches attempt to learn unified representations across many datasets^8^, yet state-of-the-art foundation models still exhibit substantial residual batch effects, reflecting persistent difficulty in distinguishing technical variation from true biological signal. Thus, new computational strategies are needed to globally remove batch effects to build universal representations of cell states and state-transitions.

In addition, a major challenge for global perturbation classification arises from the strong context dependence of perturbation effects^12^. The same perturbation can elicit markedly different transcriptional responses across cell types or environmental conditions, making it difficult to compare perturbations or predict perturbation responses using measurements acquired in different contexts. Moreover, different perturbation datasets often probe non-overlapping or partially overlapping sets of perturbations, and it is challenging to generate embeddings that allows different perturbations to be assessed in a universal representation. These limitations highlight the need for computational approaches that can disentangle the intrinsic properties of perturbations from the contexts in which they are measured, thereby enabling a global classification of perturbations.

In this work, we developed a new framework for generating a global cell-state manifold, systematically understanding cell-state transitions on this manifold, and improving prediction accuracy of perturbation responses. We first introduced the Single Cell Manifold Generator (SCMG), a contrastive learning framework that combines the advantages of pairwise dataset integration and deep learning to enable global-scale cell-state embedding and effective batch-effect removal. Using SCMG, we constructed a unified reference map that integrates most human and mouse cell types into a coherent and biologically meaningful manifold. Through a global analysis of the cell-state space, in conjunction of perturbation data analysis, we identified gene-expression programs that define major axes of cell-state variations. Building on this foundation, we developed a deep-learning model to generate universal, context-independent functional embeddings of genetic perturbations, allowing us to derive major programs/classes of perturbations and improve prediction accuracy of perturbation responses. Using this framework, we systematically analyzed several Perturb-seq datasets across different cell types and uncovered both conserved patterns and context-specific rewiring of perturbation responses. To further validate the generalizability of our framework and broaden our understanding of perturbation responses, we performed a genome-scale CRISPRi Perturb-seq screen in human embryonic stem cells (hESCs). Analysis of this new dataset not only validated the generalizability of the above-derived, conserved perturbation-response patterns, but also revealed distinct classes of differentiations in hESCs, including early germ-layer differentiations and stress-induced mesenchymal transitions.

## Results

### Construction of a global manifold of cell states

Here, we developed a contrastive learning method, SCMG, that embeds single-cell transcriptomes across diverse datasets into a 512-dimensional latent space (**Figure 1A**). We trained the SCMG with two complementary inputs: (i) transcriptome-wide expression vectors from a large number of scRNA-seq datasets and (ii) graphs derived from pairwise integrations between datasets that share overlapping cell types (**Methods**). In SCMG, an encoder maps expression vectors into the latent space, while a decoder (conditioned on dataset identity) reconstructs gene-expression profiles. A reconstruction loss (mean squared error between input and decoded expression vectors) is used to minimize information loss during dimensionality reduction. In addition, for contrastive learning, we selected pairs of datasets containing overlapping cell types, performed integration of the common cell types in each dataset pair, and connected cells belonging to biologically similar states with edges (**Methods**). We then included a contrastive loss term in SCMG to enforce cells connected by edges, including those from different datasets, to be closer than unconnected cells in the same dataset, reducing batch-to-batch variations in the embedding space. We reason that such pairwise integration overcomes the challenges associated with different annotation taxonomies and granularities across datasets and additionally allows fine-scale gene-expression variations within cell types to be aligned across datasets, which should improve the efficiency of batch correction. By minimizing both the reconstruction and contrastive loss terms simultaneously, SCMG seeks to learn a general strategy to remove batch effects while preserving biological variations.

**Figure 1.**
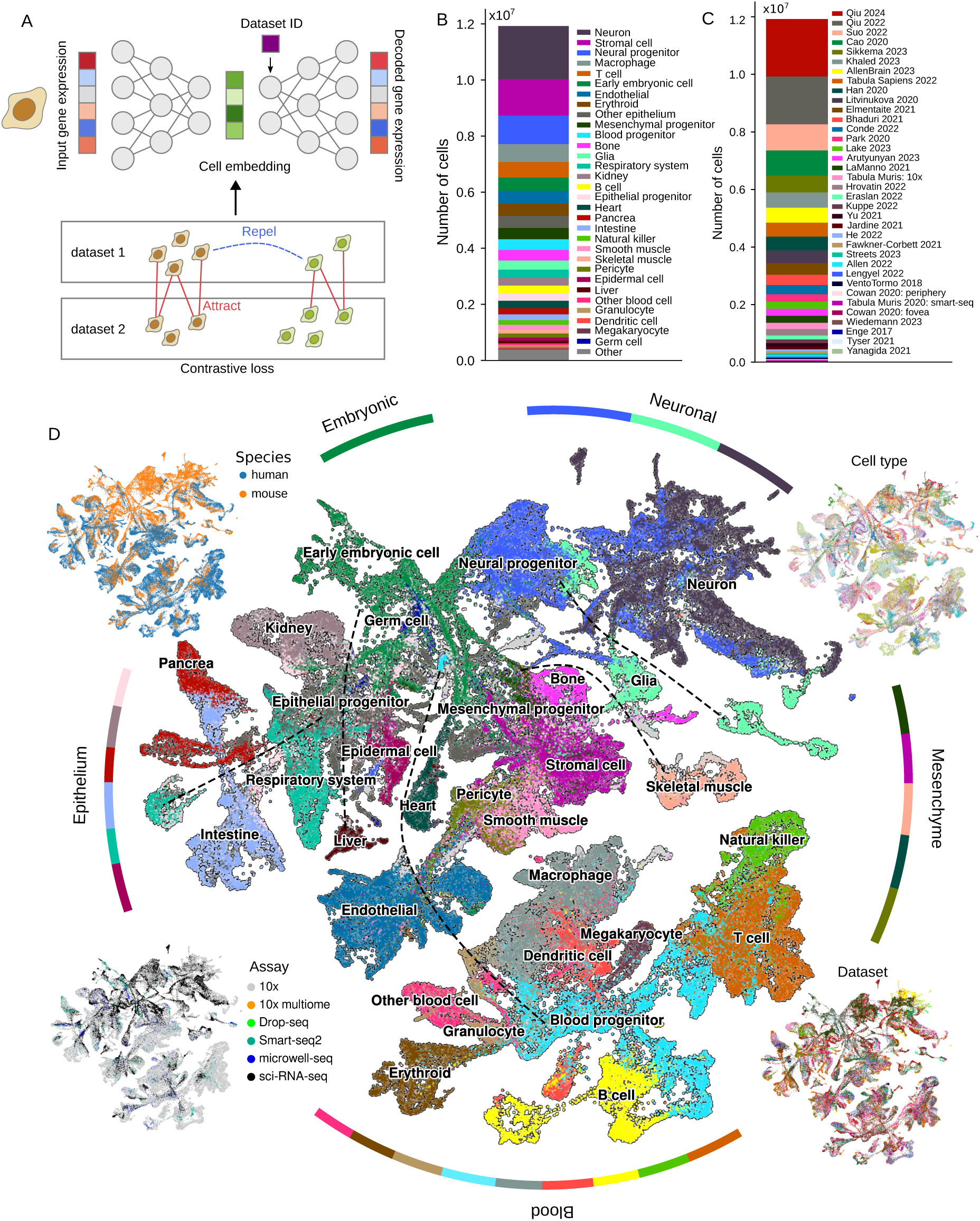
| Construction of a global cell-state manifold by a contrastive learning model. **A,** Architecture of the single-cell manifold generator (SCMG) model. scRNA-seq profiles are mapped to a 512-dimensional latent space with an encoder–decoder minimizing reconstruction loss. A graph-based contrastive objective, derived from pairwise dataset integrations, pulls together biologically matched cells (edge-connected) and separates unmatched cells, thereby reducing batch effects. **B–C,** Composition of scRNA-seq compendium, by major cell types (B) and by source datasets (C), used for training the SCMG. The compendium includes 12 million cells from 36 datasets. **D,** UMAP visualization of the global cell-state map generated by SCMG. For visualization purpose, datasets are down-sampled to provide a balanced representation across cell types. The central panel is colored by major cell type. Surrounding panels show the same UMAP colored by species (upper left), original dataset cell type annotations (upper right), assay (lower left), and origin of dataset (lower right). Dash lines connect cell types that are adjacent to each other in the 512-dimensional latent space but separated in the 2D UMAP.

Applying SCMG to 36 scRNA-seq datasets (∼12 million cells) spanning diverse human and mouse cell types^13–46^ (**Figure 1B, C**), we constructed a global manifold of human and mouse cells (**Figure 1D**). Given the extensive conservation of cellular states across these species, as well as the widespread use of mouse models to study human biology, we prioritized shared structure by preserving cell-state variations common to both human and mouse while treating species identity as batch effects. This design enables the construction of an interpretable, global reference map of conserved cell states, facilitating cross-dataset and cross-species comparisons. At the same time, we note that this approach may attenuate genuine species-specific differences in certain contexts, and our analyses are therefore focused on the conserved dimensions of cell-state organization captured by the model.

To visualize the global organization of the cell-state landscape, we down-sampled the training compendium of scRNA-seq datasets to a “representation dataset” with balanced cell-type coverage (**Methods**) and projected cells onto a two-dimensional UMAP^47^ (**Figure 1D**). In this embedding, distinct cell types segregate cleanly, whereas similar cell types from different datasets, assays, or species co-localize. Because early embryonic and neuronal cells are predominantly from mouse studies, the corresponding regions are mouse-enriched. The map recapitulates broad developmental lineages while resolving fine-grained structure. From embryonic stem cells, trajectories diverge into three major branches aligned with the three germ layers: ectoderm, mesoderm, and endoderm. The neural ectoderm branch contains neural progenitors, neurons, and glia. A mesenchymal branch, largely mesodermal, comprises mesenchymal progenitors, bone cells, stromal cells, smooth muscle cells, endothelial cells, and pericytes. Hematopoietic cells, also mesodermal, form a tree-like structure radiating from blood progenitors into distinct lineages of blood and immune cells. Most endoderm-derived cells occupy an epithelial branch, including epithelial progenitors, intestinal cells, pancreatic cells, and cells from the respiratory system. Some ectodermal (epidermis) and mesodermal (kidney) cell types also reside within the epithelial branch, likely reflecting convergent activation of shared programs (e.g., cell-cell junctions). Beyond global organization, the manifold resolves subtype-level distinctions; for example, known endothelial subtypes^48^ are well separated (**Extended Data Figure 1**).

### Understanding cell states and state transitions with the global cell-state manifold

Having qualitatively demonstrated SCMG’s ability to generate an interpretable global landscape of cell states, we next quantitatively benchmarked its performance—and that of the resulting cell-state map—across several key analytical tasks. Specifically, we evaluated three capabilities: (1) zero-shot cross-dataset integration, (2) reconstruction of gene-expression profiles from the latent space, and (3) projection of new datasets onto the global reference cell-state manifold.

First, we quantified the generalizability of SCMG’s batch correction via zero-shot integration. We evaluated co-embedding of ten dataset pairs. Among the ten, five pairs each included one dataset withheld from SCMG training. Each pair contained overlapping cell types, and we restricted analysis to those shared cell types so that successful integration would yield strong intermixing between datasets while maintaining separation of distinct cell types. We compared SCMG with five state-of-the-art deep-learning models trained on large compendia of scRNA-seq data: scVI^49^, Geneformer^50^, scGPT^51^, SCimilarity^52^, and UCE^53^. UMAPs of the co-embedded cells showed that SCMG achieved better intermixing than the other five models (**Figure 2A**).

**Figure 2.**
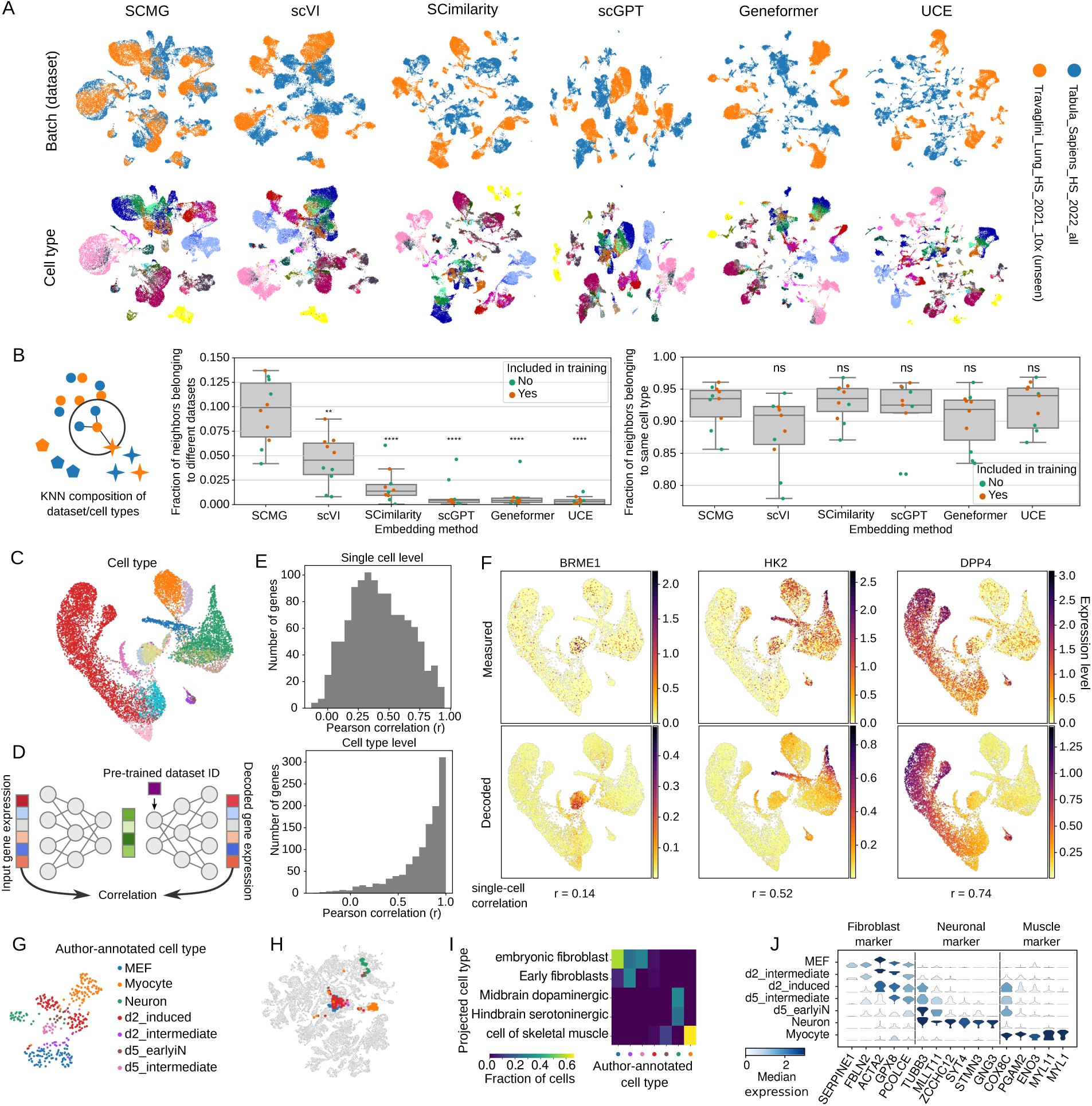
| Zero-shot integration, gene-expression decoding, and cell-state projection by SCMG. **A,** UMAP visualization of co-embeddings of a representative dataset pair using different deep-learning models (columns). Cells are colored by dataset (top) or by cell type (bottom). The Travaglini_Lung_HS_2021_10x dataset was withheld from training. Only cells with shared cell-type labels between the two datasets are shown. **B,** Quantification of integration performance based on the composition of cell types and datasets among each cell’s k nearest neighbors in the embedding space. In the schematic (left), datasets are indicated by color and cell types by shape. For each dataset pair, a k-nearest-neighbor graph (k = 30) is built with each nearest-neighbor cell pair connected by an edge, and an edge is classified as an inter-dataset edge if the two cells are from different datasets and a same-cell-type edge if the two cells belong to matching cell types. Box plots show the fraction of all nearest-neighbor edges in the graph that are inter-dataset neighbors (middle) and same-cell-type neighbors (right) for different deep-learning models. Each data point represents a dataset pair with colors indicating whether both datasets in the pair were seen during SCMG training. ** represents p-value < 0.01; **** represents p-value < 10^-4^; ns represents p-value > 0.05. **C,** UMAP of intestinal cells from Ref. 54. a dataset unseen during SCMG training. Cells are colored by author-provided cell-type annotations. **D,** Schematic of the decoding evaluation with Pearson correlation between originally measured and SCMG-decoded expression. **E,** Distributions of Pearson correlation coefficients between originally measured and SCMG-decoded expression values for individual genes across cells (top) and across cell types (bottom). **F,** Examples of measured (top) and decoded (bottom) expression for three representative genes visualized on UMAP. Correlation coefficients between measured and SCMG-decoded expression levels across cells are shown below the decoded panels. **G**, UMAP of cells from a mouse embryonic fibroblast (MEF) trans-differentiation dataset^56^, colored by author-annotated cell types. **H,** Projection of cells from the MEF trans-differentiation dataset onto the global cell-state manifold, colored by author-annotated cell types as in (**G**). **I**, Confusion matrix comparing author-annotated cell types, colored as in (**G**), with SCMG-projected cell types. **J**, Violin plots of marker gene expression across author-annotated cell types. The physiological cell types in which each gene is normally expressed are indicated above the plots.

To quantify performance, we constructed a k-nearest-neighbor graph in the latent space and computed two metrics: (i) the fraction of inter-dataset neighbor pairs (assessing batch-effect removal) and (ii) the fraction of same-cell-type neighbor pairs (assessing preservation of biological structure) (**Methods**). Compared to the other models, SCMG resulted in a substantially higher inter-dataset neighbor fraction without significantly reducing the same-cell-type fraction (**Figure 2B**), indicating more effective batch correction without losing true biological variations. Notably, SCMG’s performance was comparable for dataset pairs composed entirely of training datasets and those including an unseen dataset, demonstrating efficient zero-shot integration. Among the methods, those that explicitly correct batch effects (SCMG, scVI, SCimilarity) outperformed models primarily optimized for gene-gene relationships and transfer learning. SCMG further surpassed scVI and SCimilarity in zero-shot integration, likely because the training input in SCMG includes supervision from pairwise dataset integrations, which could provide a stronger and more faithful signal for disentangling batch effects from true biological variation than either unsupervised correction (scVI) or discrete, pre-annotated cell-type labels (SCimilarity), which may have inconsistencies between datasets and may suppress continuous cell state variation within the same cell type.

Second, we quantitatively assessed SCMG’s preservation of biological variation by comparing measured expression with decoder reconstructions in datasets unseen during training (**Figure 2C-F and Extended Data Figure 2A-C**). We performed this analysis on two unseen datasets profiling the intestine and the lung^54,55^. For each gene, we computed Pearson correlations between observed and SCMG-decoded expression levels across cells and, separately, across cell types (**Methods**). Single-cell correlations spanned a broad range, whereas cell-type-level correlations clustered near one (**Figures 2E and Extended Data Figure 2B**), indicating strong preservation of cell-type-dependent gene expression patterns. At single-cell resolution, SCMG attenuated stochastic fluctuations of gene expression, yielding smoother expression profiles (**Figure 2F and Extended Data Figure 2C**). We note that while this denoising likely reduces technical noise (e.g., dropout), it may also obscure some genuine fine-scale variability.

Third, we tested SCMG’s ability to project cells from unseen datasets onto the global reference cell-state manifold (**Extended Data Figure 2D-G**). Such projection allows us to identify each query cell’s nearest cell state on the reference map (**Methods**), which enables cell-type annotation of the query cells and interpretation of their relationships to other known cell states. We projected cells from the two unseen datasets profiling the intestine and the lung^54,55^ onto the global manifold and assigned cell-type labels to these cells based on the projection. The resulting labels closely matched the authors’ original annotations, but with less granular splitting of subtypes in some cases. These results demonstrate the ability of SCMG to annotate cells from scRNA-seq datasets without retraining or fine-tuning.

Projection of cells onto the global cell-state manifold can also provide insights into how perturbations reshape cellular identities. To illustrate this, we examined a published dataset in which mouse embryonic fibroblasts were transdifferentiated into muscle cells and neurons through transcription factor (TF) overexpression^56^ (**Figure 2G-J**). Projection onto the global cell-state manifold correctly mapped the starting population and the fully transdifferentiated cells to embryonic fibroblasts, mid–hindbrain neurons, and skeletal muscle cells, respectively (**Figure 2H, I**). Interestingly, intermediate states along both differentiation trajectories—particularly d2_induced and d5_earlyiN—did not consistently map to any reference cell type (**Figure 2I**). These intermediates co-expressed fibroblast, neuronal, and muscle markers (**Figure 2J**), including markers such as PCOLCE, TUBB3, and COX8C that are normally mutually exclusive across physiological cell types, suggesting that TF-induced transdifferentiation temporarily moves cells out of the physiological cell-state manifold, with cells re-entering the manifold once transdifferentiation is complete.

### Global identification of gene-expression programs

Having constructed a global reference map of cell states, we next sought to define a principal set of axes of cell-state transitions that substantially reduce the dimensionality from thousands of genes to a tractable and interpretable number while capturing the dominant variations across the global cell-state landscape. With this dimensionality reduction, we will be able to systematically interpret how a perturbation causes cells to change along each of the principal axes and facilitate comparison of perturbation responses from different datasets. We defined these principal axes by identifying the co-regulated gene expression programs, i.e. a set of genes that co-vary across the global cell-state space.

Ideally, we would derive gene-expression programs from a comprehensive set of human datasets measured with a single assay and spanning the full cell-state landscape. In practice, the available human datasets are still incomplete—especially for embryonic stages—so mouse datasets are needed to fill the gaps, which would introduce species-specific expression differences. Assay heterogeneity poses a second challenge: although most cells are profiled with the 10x scRNA-seq platform, embryonic regions of the landscape are dominated by data generated using sci-RNA-seq. Although SCMG could remove such assay-specific batch effect, and possibly some species-specific differences in the latent space, these differences will be recovered in the decoded gene expression vectors when the original dataset IDs are used as input. To remove such differences and recover evolutionarily conserved programs, we reconstructed each cell’s expression profile using the SCMG decoder, but conditioning on the dataset ID of the nearest human cell profiled with the 10x assay (**Figure 3A**, **Methods**). This effectively converts every single-cell transcriptome profile into a “human/10x-equivalent” expression vector, suppressing assay- and species-specific variation and allowing downstream analyses to focus on conserved expression patterns.

**Figure 3.**
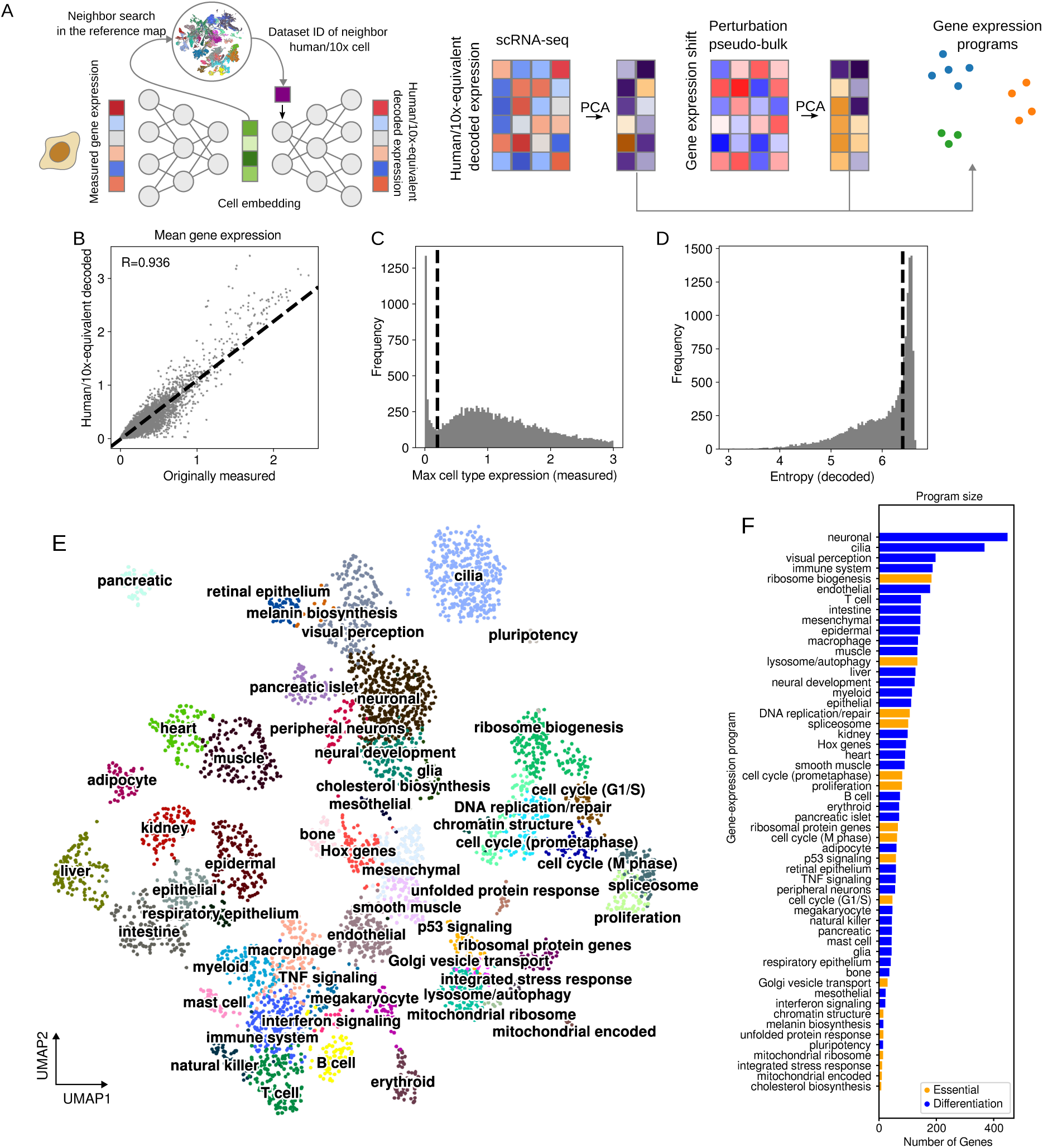
| Systematic identification of gene-expression programs to define major axes of cell-state transitions. **A,** Schematic of the workflow. Left: gene expression profiles are encoded into the SCMG latent space and decoded with dataset IDs of the nearest-neighbor “human/10x” cells to remove species/assay effects. Right: Gene expression programs are inferred from the global cell-state reference map (using “human/10x-equivalent” decoded expression vectors) and the standardize perturbation database (as described in **Figure 2K**) by PCA-based dimension reduction followed Leiden clustering. **B,** Scatter plot comparing the mean measured versus “human/10x-equivalent” decoded expression averaged across all cells the global cell-state reference map. Each dot represents a gene. The Pearson correlation coefficient r is indicated. **C,** Distribution of the peak expression of individual genes (namely, the expression level of the gene in its highest-expression cell type) after “human/10x-equivalent” decoding. **D,** Entropy of cell-type-level gene expression profiles used to filter out genes that exhibit nearly uniform expression across the cell-state map. The expression profile of each gene across the global cell-state manifold is normalized and treated as a probability distribution, and the entropy of the distribution is computed for the gene, with high entropy values representing gene with more uniform expression across the manifold (**Methods**). **E,** UMAP visualization of the Leiden clustering of genes to derive the gene-expression programs. Genes are colored by their program labels. We note that some Leiden clusters lacked clear cell-type specificity or gene-set enrichment, and hence are unannotated and not included in (E) and (F) and subsequent analysis. **F,** Number of genes in each gene-expression program. “Essential” and “differentiation” programs are colored in orange and blue, respectively.

The human/10x-equivalent decoded and measured expressions of individual genes showed strong correlation (Pearson r = 0.936, **Figure 3B**). Using the human/10x-equivalent expression profiles, we sought to identify gene-expression programs, which are defined as clusters of genes that share similar expression patterns across the cell-state manifold (i.e. genes that co-vary across cell states). Because our global cell-state manifold spans most mammalian cell types, it is well-suited to recover differentiation-related gene programs. However, as the compendium of scRNA-seq datasets largely comprises physiologic samples, stress-response programs could be underrepresented. To obtain a more comprehensive set of gene programs, we gathered nine datasets of RNA-seq measurements before and after genetic perturbations (knockdown/knockout or overexpression) to generate a perturbation database, including both single-cell and bulk measurements of mouse or human cells, covering perturbations of 7270 genes in total (**Method**s)^57–63^. We jointly analyzed the physiological cell-state manifold with the genetic perturbation datasets (**Figure 3A**, **Methods**), using the following approach.

First, we filtered genes to focus program identification on genes that (i) vary meaningfully across cell states or perturbation conditions and (ii) participate in coherent co-variation structures, rather than exhibiting isolated or near-uniform patterns. We performed this filtering separately for physiological and perturbed conditions (**Methods**). For the physiological conditions on the reference cell-state manifold, we excluded 2,376 genes with low expression in all cell types (**Figure 3C**). We then quantified how uniformly each gene is expressed across the cell-state manifold using an entropy-based approach, with high entropy values meaning high uniformity of expression across cell states (**Figure 3D**). To remove these uniformly expressed genes that are uninformative to cell-state transitions, we selected an entropy cutoff that excluded 6,451 genes, leaving 9,281 genes that were differentially expressed across cell states.

Next, we defined each gene’s expression vector as its expression profile across all single cells in the reference cell-state manifold and computed Pearson correlation coefficients, *r*, between all pairs of genes. For each gene, we counted the number of highly correlated partners (*r* > 0.5) and retained only those genes that have at least 5 such partners and hence likely belong to a gene-expression program, yielding 8,544 genes. Finally, we performed principal component analysis (PCA) to reduce the dimensionality of the gene-expression vectors to 50 (**Figure 3A**). Applying a similar approach to the perturbation database, we identified 1,613 readout genes that exhibited strong perturbation responses and had co-varying partners. We then computed pseudo-bulk expression-shift vectors (perturbation-induced changes in expression) for these genes across perturbations and reduced the dimensionality of these perturbation-shift vectors to 50 for each readout gene (**Figure 3A**).

From the union of genes selected in the physiological and perturbation analyses (9,345 genes total), we concatenated each gene’s physiological and perturbation PCA embeddings to form a 100-dimensional feature vector. We then applied Leiden clustering^64^ in this joint space to identify gene-expression programs (**Figure 3A**, **Methods**), yielding 54 programs comprising 8 to 448 genes each (**Figure 3E, F**). We annotated these gene programs using their mean expression profiles on the global cell-state manifold and/or their enrichment of previously curated gene sets (Gene Ontology^65^, Reactome^66^, CORUM^67^ and KEGG^68^), and group these programs into two classes, “essential” and “differentiation”, based on their expression patterns and the functions of their member genes.

The “essential” class contains several programs associated with core cellular processes, including chromatin structure, DNA replication/repair, cell-cycle control, RNA splicing, and ribosome biogenesis (**Figure 3E**). Their expression on the global cell-state manifold highlights proliferative cell populations—most prominently early embryonic cells and hematopoietic progenitors (**Extended Data Figure 3**). In contrast, most mature lineages downregulate (but still express) these programs after development, with the exception of specific stem and progenitor populations which showed relative high expression of these programs. Although these programs all contribute to proliferation, the fact that their constituent genes partition into distinct programs suggests that they are regulated by separable mechanisms. Several other “essential” programs—such as ribosomal protein genes, mitochondrial encoded genes, Golgi vesicle transport, unfolded protein response, integrated stress response, p53 signaling, and lysosome/autophagy—are broadly expressed on the physiological cell-state manifold (**Extended Data Figure 3**), indicating that they vary more strongly in response to perturbations than across cell types. Notably, the cholesterol biosynthesis module, which we classified as an “essential” program because cholesterol is required in all human cells, showed pronounced cell-type variability, with glia, immature neurons, and liver cells exhibiting the highest expression levels (**Extended Data Figure 3**). This pattern is consistent with prior studies demonstrating elevated cholesterol biosynthesis activity in these cell types^69–71^.

The “differentiation” class comprises programs that span multiple levels of cell-type specification (**Figure 3E**). At the broadest scale, each major lineage—embryonic, neuronal, mesenchymal, epithelial, and hematopoietic/immune—maps to a distinct program. Most major cell types within these lineages also have their own program. These observations suggest that at coarse granularity, cell-type identities are defined by strongly correlated transcriptional modules, whereas finer distinctions arise from a smaller set of specific genes or combinatorial expression patterns. We also identified several cross-lineage programs (**Extended Data Figure 3**). For example, a HOX-gene-enriched program is active across most progenitor populations (except for the blood cells), consistent with its conserved role in developmental patterning^72^. A large cilia program is strongly induced in ciliated nodal cells (embryonic), ciliated epithelial cells, and ependymal cells (neuronal), reflecting the well-known multiciliated cell types^73^. Similarly, interferon and TNF-signaling programs are expressed across mature immune, mesenchymal, and epithelial cell types under physiological conditions, consistent with the diverse cell types that can activate these signaling pathways^74,75^.

Overall, the gene expression programs that we derived from the global cell-state manifold and perturbation database provide a principal decomposition of high-dimensional transcriptomes into a reduced number of biologically interpretable axes that can help understand cell-state determination and transition.

### Global identification of perturbation classes

Having established a global set of gene-expression programs to enable systematic descriptions of cell-state variations across datasets, we next turned to the perturbation space and sought a global representation and classification of genetic perturbations. As is for the gene expression space, the space of genetic perturbations is vast. Grouping these perturbations into a smaller number of interpretable classes would both clarify the structure of gene-function space and provide a systematic way to compare how perturbation responses (and hence functional pathways) manifest—and rewire—across different cell contexts (e.g. cell states or environmental conditions).

Perturbation-perturbation correlations have often been used to identify and group genes with similar functional impact into modules. However, a key challenge remaining in perturbation-based classification of genes is that perturbation-induced transcriptional shifts are strongly dependent on the cell context, such as the initial cell state. To illustrate this, we computed cosine similarities between gene-expression shift vectors induced by the same perturbation from different initial cell states. Plotting these similarity scores against both the distances between initial cell states on the global cell-state manifold and the perturbation-induced gene-expression shift magnitudes showed a strong dependence of the perturbation effects on the initial cell state (**Figure 4A**). This dependency indicates that directly comparing shift vectors across contexts conflates intrinsic gene function with context-specific response programs. This motivates the development of a method that explicitly disentangle intrinsic functional effects from context-dependent responses in perturbation data.

**Figure 4.**
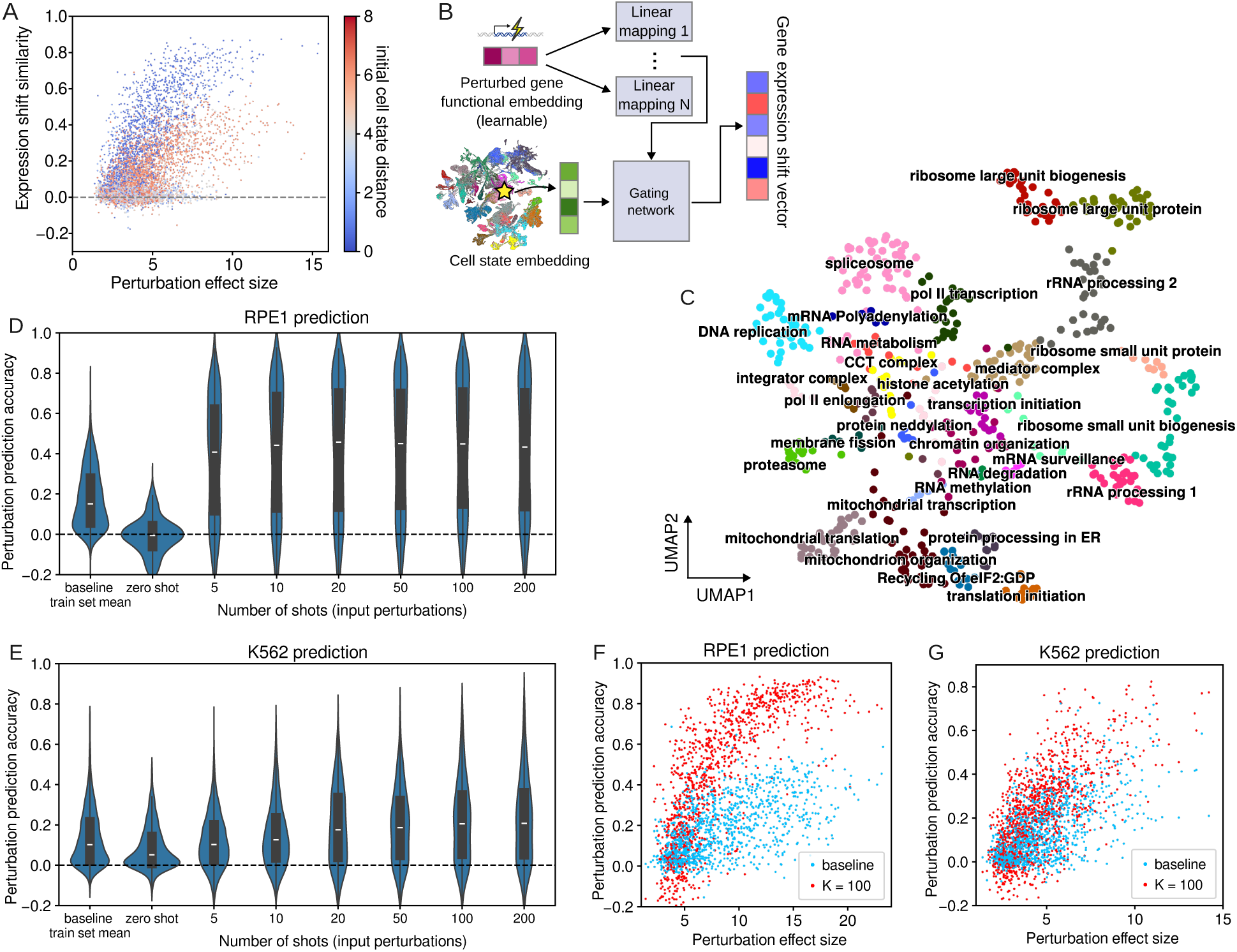
| Global functional embeddings of genetic perturbations and few-shot prediction of perturbation responses. **A**, Cell-state dependence of perturbation response. To compare the gene-expression shifts induced by the same perturbation from two different initial cell states, we quantified the difference of the two initial state by their distances in the 512-dimensional latent space and perturbation response similarity from these two states as the Cosine-similarity between the two expression shift vectors (y axis). The plot shows the dependence of the perturbation-induced gene-expression shift similarity (y axis) on the initial cell-state distance (color) and perturbation effect magnitude (x axis, represented by the smaller of the magnitudes of the two gene-expression shift vectors). Each point represents a perturbation. **B,** Schematic of the approach to learn functional embeddings of the perturbed genes. A mixture-of-experts maps functional space to gene-expression shift vectors conditioned on initial cell state. The model assumes a universal embedding space of perturbed gene functions and that the functional embedding space can be linearly mapped to perturbation-induced gene expression shifts under each cell-state context. Different cell states have different linear maps which are linear combinations of N basis linear mapping functions. The coefficients of the linear combination are determined by a gating network that conditions on the initial cell states. Training of the mixture-of-expert model learns the universal functional embedding from diverse perturbation datasets. **C,** UMAP visualization of Leiden clustering of the learned functional embeddings for genes. Only genes with robust and reproducible measured effects are included (filtered as described in the main text). Points are colored by annotated functional clusters. **D, E,** Violin plots showing the distribution of prediction accuracy for RPE1 (**D**) and K562 (**E**) datasets. The prediction accuracy is measured by Pearson correlation between the true perturbation-induced expression shift vector (measured by Perturb-seq) and the predicted shift. Each violin shows the results from a few-shot model trained with the specified number of shots (K), along with a zero-shot model and a simple baseline model (as described in the main text). The value of the shot number K corresponds to the number of perturbed genes, whose measured perturbation responses are included in the few-shot training. **F, G,** Scatter plot showing the perturbation prediction accuracy (as described in D and E) as a function of measured perturbation magnitude using the few-shot framework with K = 100 (Red) and the simple baseline model (Cyan). (**F**) For RPE cell dataset. (**G**) K562 cell datasets.

We leveraged SCMG’s global cell-state embedding as a continuous descriptor of cell-state context to separate intrinsic gene function from cell-state-specific perturbation responses (**Figure 4B**). Specifically, we posit a global latent functional space in which functionally similar genes (i.e. genes whose perturbations elicit similar cellular responses) occupy nearby positions in this space. The transcriptional response of perturbing a gene can then be obtained by mapping its functional embedding to a perturbation-induced expression shift vector through a transformation that depends on the starting cell state. To implement this framework, we developed a mixture-of-experts model^76^: each expert is a linear mapping from functional embeddings of perturbed genes to perturbation-induced expression shift vectors, while a gating network conditioned on the SCMG cell-state embedding assigns context-dependent weights to the experts. The model’s predicted response (expression shift vector) is then a weighted combination of the experts’ outputs (**Methods**). Because the functional embedding is shared across all training data and thus context independent, this architecture supports joint learning across multiple perturbation datasets—even when different experiments perturb different subsets of genes—while explicitly modeling context dependence through the gating network.

We trained the mixture-of-experts model on seven published Perturb-seq knockdown datasets^57–59,61,62^, three of which contain genome-scale perturbations (on K562 and RPE1 cells) and the rest at a smaller scale, and used the resulting functional embeddings to organize perturbations by similarity. Although the model can, in principle, incorporate arbitrarily many datasets, publicly available Perturb-seq collections currently cover a limited number of cell-type contexts. In addition, many perturbations produce only weak transcriptional shifts, yielding low signal-to-noise ratios and limited reproducibility across biological replicates. To ensure that downstream clustering is driven by reproducible perturbation signatures and thus reduce noise-driven false discoveries, we restricted downstream analyses to 608 genes, the perturbations of which led to pseudo-bulk expression-shift vectors that were concordant between two independent genome-scale Perturb-seq experiments in K562 cells (Pearson correlation > 0.4). Applying Leiden clustering to the functional embeddings of these genes, and annotating the resulting clusters using enriched Gene Ontology terms, revealed diverse cellular pathways spanning processes from the central dogma to mitochondrial function (**Figure 4C**). Together, these results suggest that the learned functional embedding space captures coherent, biologically interpretable structure across perturbations.

A global embedding of perturbed gene functions also enables few-shot prediction of perturbation responses. Because genes that elicit similar transcriptional shifts when perturbed are placed near one another in the embedding space, measured responses for a small and representative set of perturbed genes can inform perturbation responses of nearby genes. If the functional embedding captures intrinsic relationships among gene functions that extend beyond the specific contexts (i.e. initial cell state) seen during training, we reason it is then possible to use this functional embedding to improve the prediction accuracy of perturbation responses of essentially all genes in a different context by profiling, in that context, perturbations of only a limited set of representative genes that efficiently sample the functional embedding space. The strong correspondence between the learned embedding structure and canonical molecular pathways (**Figure 4C**) supports this hypothesis.

To test this idea of few-shot prediction, we retrained two separate mixture-of-experts models while withholding, from the seven training perturb-seq dataset, perturbation experiments performed in K562 and RPE1 cells, respectively, and evaluated few-shot prediction on each held-out dataset. Specifically, for each held-out dataset, we selected K genes (K = 5–200) that sampled the functional embedding space and fit linear models that map functional embeddings to the measured perturbation-induced expression-shift vectors for these K genes (**Methods**). We then used these models to predict the perturbation responses of the other genes and assessed the prediction accuracy as the Pearson correlation between predicted and experimentally measured expression shift vectors. As a comparison, we also assess the prediction accuracies of two other approaches: (i) a zero-shot model by directly using the mixture-of-experts model trained as described in **Figures 4B**, without any input from the holdout dataset; (ii) a simple baseline model that predicts the perturbation-induced expression-shift vector of a gene knockdown as the mean value of shift vectors with the same perturbation across the training dataset. This latter baseline model was previously used to shown that previously existing deep-learning approaches do not output simple linear baselines in prediction accuracy^9^. Notable, our few-shot models substantially outperformed both this baseline model and the Zero-shot model—even with as few as K = 10 training perturbations (perturbed genes) (**Figures 4D,E**). Accuracy increased with larger K and plateaued around K ∼ 100, and predictions were more accurate for genes with larger perturbation effects, consistent with improved signal-to-noise ratios (**Figures 4F,G**). Few-shot prediction performed better in RPE1 cells, potentially reflecting convergence of diverse perturbations onto a smaller set of transcriptional phenotypes in this context.

Although the absolute prediction accuracy remains imperfect given the limited size of available training data, these results provide a proof of principle that interpretable, global functional embeddings obtained using our approach can support few-shot perturbation prediction and improve the accuracy of perturbation prediction. It is interesting to note that while zero-shot prediction performance is worse than the baseline, the few-shot prediction performance surpassed the baseline, potentially because the functional pathway relationship is less context-dependent and therefore easier to learn, whereas the specific perturbation effects is context-dependent and hence more difficult to learn from the limited existing data.

### Conservation patterns and cell-state-specific rewiring of perturbation effects

Having established a global framework for representing cell states, state transitions, and perturbations, we next used this framework to compare perturbation-induced cellular responses across several Perturb-seq datasets acquired in different cell-state contexts. We focused on two questions: (i) to what extent do perturbations of essential cellular processes couple to differentiation programs, and (ii) how do functional pathways rewire across cell states, altering their transcriptional impact in a context-dependent manner?

Essential cellular processes and differentiation are tightly linked, at least in one direction: differentiation often entails large changes in essential processes (for example, cell-cycle exit accompanies many lineage transitions). What is less clear is how often the relationship runs in the opposite direction—whether perturbing essential cellular pathways is sufficient to engage or redirect differentiation programs. Previous studies have reached different conclusions: some argue that perturbation of essential pathways do not influence differentiation, whereas others suggest that reciprocal regulation between these processes is fundamental for development^77,78^. Synthesizing these results across studies obtained from different contexts is challenging. We therefore sought a systematic, data-driven assessment of coupling between essential processes and differentiation. To this end, we applied our framework of gene-expression programs to three large-scale perturbation screens: genome-scale CRISPR interference (CRISPRi) knockdown Perturb-seq datasets of K562 and RPE1 cells^57^, and an open reading frame (ORF) overexpression screen of human embryonic stem cells (hESCs)^63^ that primarily targets transcription factors, along with a subset of ligands and receptors. For each perturbation, we computed a pseudo-bulk gene-expression shift vector and quantified shift scores for every gene-expression program derived in this study, defined as the weighted average expression change of the program’s constituent genes with weights chosen to account for stochastic variability (**Figure 5A, Methods**).

**Figure 5.**
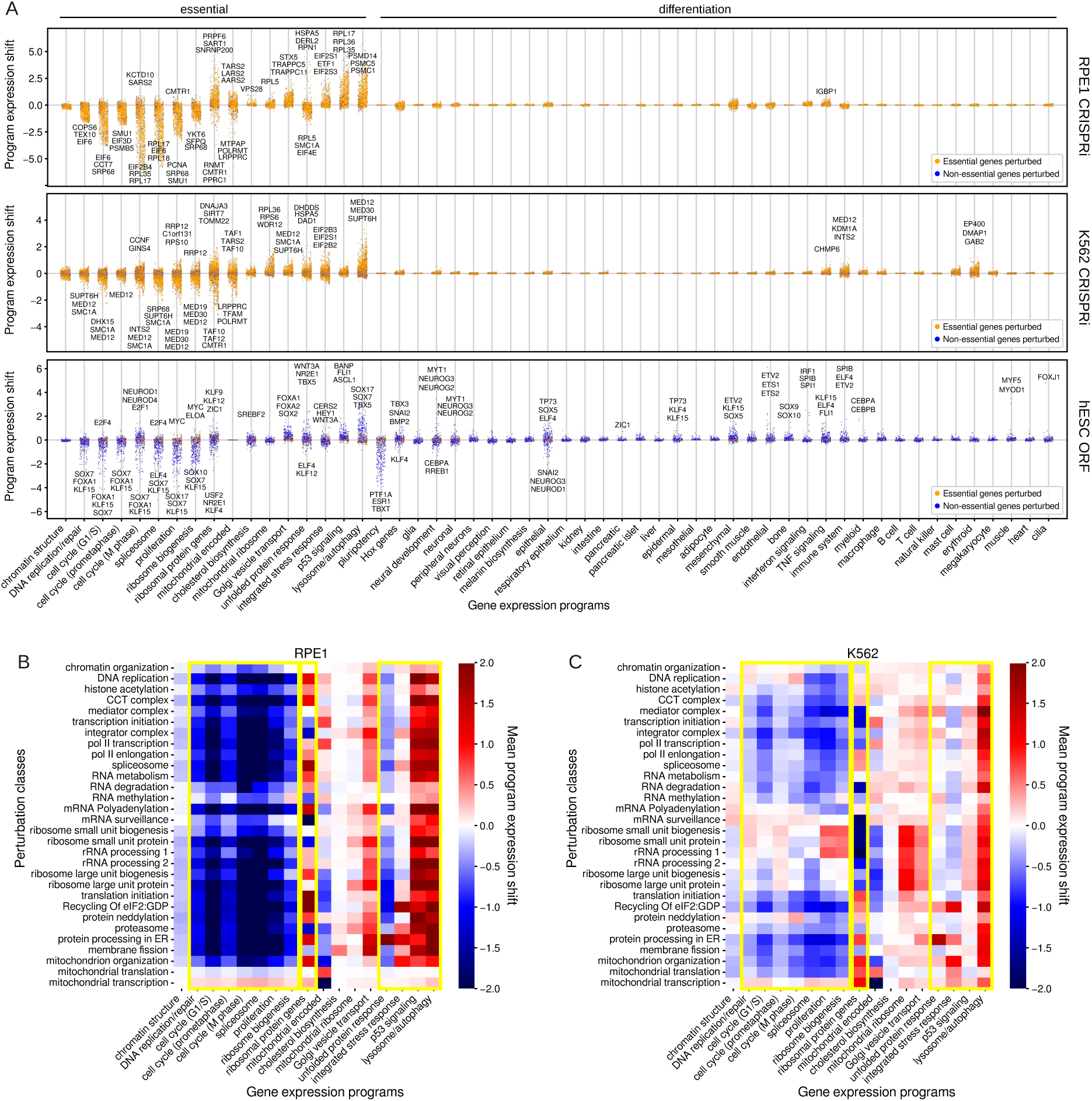
| Conserved patterns and cell-type-specific rewiring of changes in gene expression programs induced by genetic perturbations. **A,** Expression shift scores of gene-expression programs for each perturbation across three datasets: RPE CRISPRi perturb-seq (top), K562 CRISPRi perturb-seq (middle), and hESC ORF overexpression screen (bottom). The program expression shift score presented is the mean score across all perturbations within each class. Gene names of the top 3 perturbations inducing the largest shifts in each direction are labeled if their absolute program expression shift scores are greater than 1. Each point corresponds to a genetic perturbation, colored in orange and blue for essential and non-essential genes, respectively. Essential and Differentiation gene-expression programs at labeled at the top. **B, C,** Heatmap of essential program changes induced by each perturbation classes in RPE1 (**B**) and K562 (**C**). Yellow boxes highlight gene expression programs related to cell proliferation, ribosome protein genes, and stress response.

We operationally classified perturbed genes into essential and non-essential categories: a gene was considered essential if it was broadly expressed across the global cell-state map or if it belonged to one of the essential gene-expression programs. In both CRISPRi datasets, we observed that the gene-expression programs belonging to the differentiation category (as defined in **Figure 3E, F**) were largely insulated from knockdown of essential genes, whereas these knockdowns substantially impacted gene-expression programs belonging to “essential” category (**Figure 5A**). As a control, we observed that TF overexpression can indeed induce broader differentiation responses, as expected (**Figure 5A**). In contrast, CRISPRi knockdown of essential genes in K562 and RPE1 cells primarily modulated programs aligned with each cell line’s baseline identity (e.g., hematopoietic programs in K562; mesenchymal and TNF-signaling programs in RPE1). Overall, these results suggest that perturbations to essential processes broadly engage stress-related gene programs but show limited ability to trigger differentiation transitions.

Next, we leveraged the global gene-expression programs together with the perturbation classes to systematically compare how perturbations drive cell-state changes, and how these relationships rewire between two cellular contexts—K562 and RPE1 cells. By plotting the mean effect of each perturbation classes, we identified distinct perturbation-response patterns involving stress-response and proliferation-related programs (**Figures 5B, C**), which is also supported by examination of canonical marker genes of the pathways represented by the perturbation classes and gene expression programs (**Extended Data Figure 4**).

Among the gene-expression programs, four correspond to major stress-response pathways: the unfolded protein response (UPR), the integrated stress response (ISR), p53 signaling, and lysosome response/autophagy. Prior studies show that each pathway can be activated by diverse stimuli—including hypoxia, nutrient deprivation, viral infection, and oxidative stress^79–82^. Our framework allows cross-dataset analysis, providing a direct view of the stimulus spectrum and context dependence of these pathways (**Figures 5B, C**). We found that the lysosome/autophagy program acts as a general stress response in both K562 and RPE1 cells that is activated by nearly all classes of essential-gene knockdowns except for mitochondrial transcription and/or translation perturbations. p53 signaling likewise behaves as a broadly responsive pathway in RPE1 cells, whereas—as expected—no p53 activity is detected in p53-deficient K562 cells^83^. In contrast, other stress pathways show more selective activation by different perturbation classes. ISR is specifically induced by perturbations affecting eIF2:GDP recycling, protein processing in the ER, and also by perturbation of the mitochondrial organization, consistent with canonical activation mechanisms^79^. Notably, mitochondrial transcription and translation perturbations activate ISR in K562 cells but not in RPE1 cells, showing context-dependent rewiring of this pathway. This difference cannot be attributed to variation in CRISPRi efficiency, as knockdown of mitochondrial transcription and translation genes induced strong and consistent shifts in mitochondrial-encoded transcripts in both cell types. Finally, as expected, the UPR is activated almost exclusively by perturbations that disrupt ER protein processing^84^. Together, these results delineate the spectrum of activators for core stress pathways and how their engagement depends on cellular context.

In contrast to stress-response programs, which are predominantly activated by perturbations, we also identified several gene-expression programs that are broadly repressed across diverse perturbations, including those related to the cell cycle, spliceosome, proliferation, and ribosome biogenesis (**Figures 5B, C**). The inclusion of cell-cycle and proliferation programs suggests that these axes likely reflect proliferative capacity, consistent with prior findings that spliceosome and ribosome-biogenesis genes are upregulated in certain proliferative cell types^85,86^. Interestingly, the program of ribosomal protein (RP) genes shows highly distinct regulation from the ribosome-biogenesis gene program. In both K562 and RPE1 cells, RP genes are upregulated upon perturbation of processes such as translation initiation and mitochondria organization, but downregulated when components of the Integrator complex or mRNA surveillance pathways are disrupted. Moreover, RP-gene program regulation is cell-type-specific: for example, perturbing large-subunit ribosome biogenesis decreases RP-gene expression in K562 cells but increases it in RPE1 cells. Our results expand on the previous suggestion of differential regulation of RP genes and ribosome-biogenesis genes^87^, providing a broader spectrum of stimuli that separates these programs, and indicate that ribosome biogenesis is coupled to proliferative capacity, whereas RP genes respond broadly to diverse cellular stresses. Although these two pathways are often treated interchangeably in the literature^86,88,89^, our findings underscore the importance of treating these programs as mechanistically distinct when studying ribosome homeostasis.

### Genome-scale Perturb-seq on human embryonic stem cells

Using our global analysis framework together with published datasets, we obtained systems-level understandings of cellular responses to genetic perturbations. However, currently available genome-scale perturbation datasets remain limited in number and sample only a narrow subset of possible cell-state contexts. Although our analytical framework is readily extensible to additional contexts, expanding our understanding of cell-state regulation ultimately requires generating new perturbation data in previously unexplored contexts. To broaden the scope of our analysis and test the generalizability of our findings, we performed a genome-scale CRISPRi Perturb-seq experiment in human embryonic stem cells (hESCs).

We selected hESCs for two reasons. First, they represent a cell state that is developmentally and transcriptionally distinct from K562 and RPE1 cells—the two lines with previously published genome-scale CRISPRi Perturb-seq datasets—providing a test of whether regulatory principles derived from those systems extend to an early developmental context. Second, our comparison of a TF overexpression screen in hESCs with CRISPRi knockdown screens in K562 and RPE1 had revealed broader differentiation responses in the hESC overexpression screen. Performing a CRISPRi screen directly in hESCs allows us to assess whether differences in differentiation outcomes reflect intrinsic properties of the cell state itself or the types of perturbations applied.

We designed a CRISPRi library comprising 6,638 sgRNAs targeting 2,978 genes selected for putative roles in hESC growth or pluripotency marker expression (**Methods**), and profiled approximately 1.3 million perturbed cells (**Figure 6A**). Each perturbation was represented by tens to hundreds of cells (**Figure 6B**), and most sgRNAs achieved effective knockdown of their target genes, with a median expression reduction of ∼70% (**Figure 6C**). Despite this robust knockdown, perturbation-induced transcriptional effect sizes were substantially smaller in hESCs than in K562 cells, as measured by the distribution of the expression shift vector magnitude across perturbations (**Figure 6D**). This attenuation cannot be explained solely by differences in knockdown efficiency: even when restricting analysis to genes with high knockdown efficiency in hESCs, the magnitudes of the perturbation-induced transcriptional shift remained significantly lower than those observed in K562. These results indicate that hESCs are in general more resistant to perturbation-induced stress, expanding on previous findings that hESCs exhibit higher resilience to some specific stressors than differentiated cell types^90,91^ and suggesting broad resistance to diverse perturbations in the hESC state.

**Figure 6.**
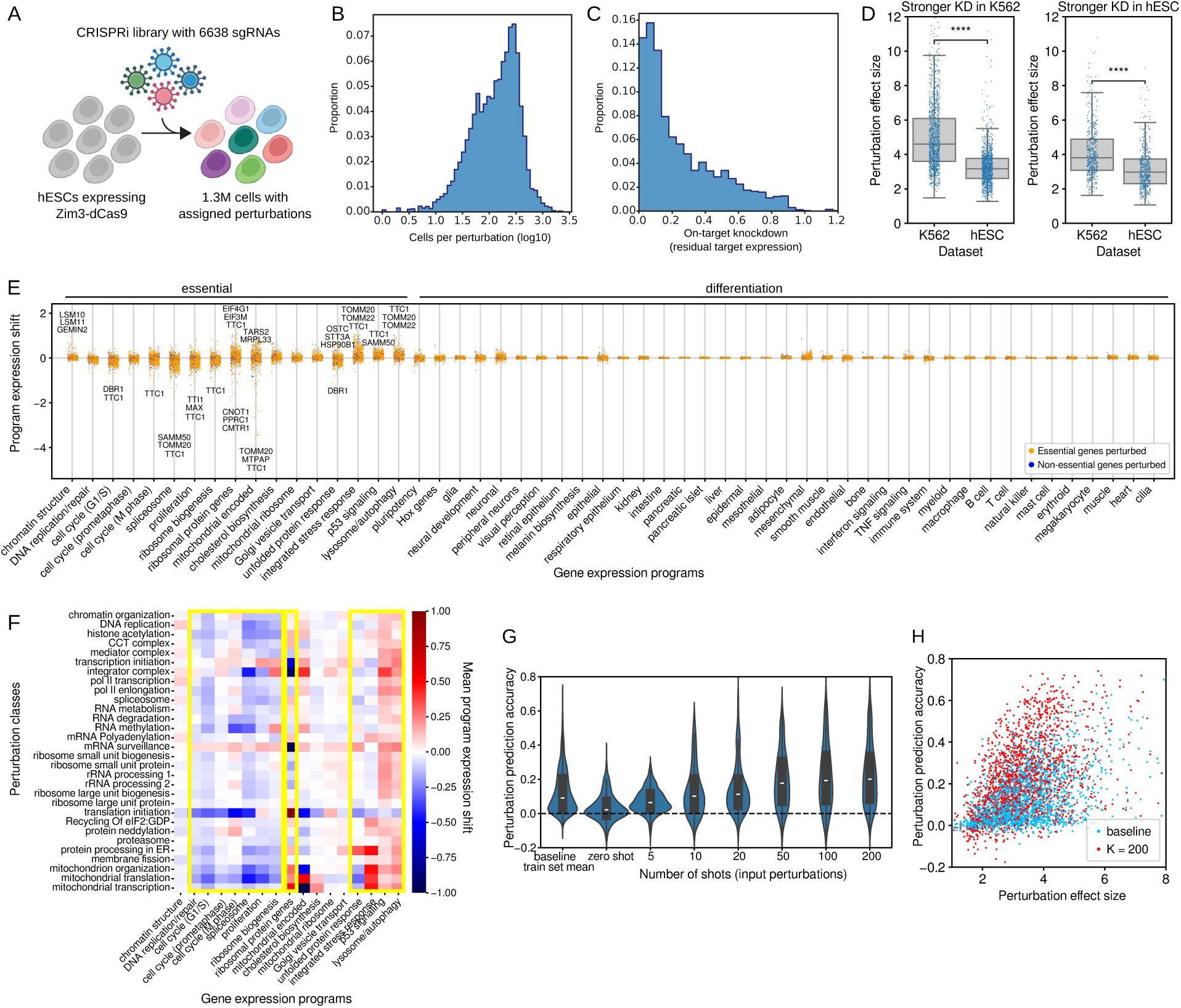
| Genome-scale CRISPRi perturb-seq experiment and pseudobulk analyses for human embryonic stem cells (hESCs). **A,** Schematic of the genome-scale CRISPRi perturb-seq screen in hESCs. **B,** Probability density distribution of the number of cells profiled per perturbation. **C,** Probability density distribution of on-target knockdown efficiencies. The normalized residue expression level is shown. **D,** The magnitude of the perturbation induced gene expression shift vectors for the hESC and K562 datasets. Left: distributions for genes with stronger knockdown efficiency in K562 cells. Right: distributions for genes with stronger knock down efficiency in hESC cells. **E,** As in **Figure 5A**, but for hESCs. **F,** As in **Figure 5B, C**, but for hESCs. **G,** As in **Figure 4D, E**, but for hESCs. **H,** As in **Figure 4F,G**, but for hESCs.

Despite the overall attenuation of perturbation effect sizes in hESCs, the structure of perturbation-response patterns that we observed in K562 and RPE1 cells is largely conserved in hESCs. First, although hESCs possess strong differentiation potential, perturbation of essential genes predominantly induced responses in essential programs with relatively small effect on differentiation programs (**Figure 6E**). Second, lysosomal/autophagy and p53 signaling pathways were broadly activated across most perturbations, whereas the ISR and UPR remained more selectively induced by specific classes of perturbations (**Figure 6F**), as in the case of K562 and RPE cells. A notable distinction in hESCs is that perturbations of mitochondrial transcription and translation elicited stress-response magnitudes comparable to or greater than those induced by other essential perturbations—contrasting the attenuated responses observed in the other two cell lines, especially in RPE1 cells. This heightened sensitivity is consistent with the established dependence of embryonic stem cell proliferation on intact mitochondrial function^92^.

Third, proliferation-associated programs showed downregulation across most perturbations, and ribosomal protein gene program continued to display regulatory behavior distinct from ribosome-biogenesis gene program, exhibiting both up- and down-regulation selectively to different responses (**Figure 6F**). Perturbations that most strongly modulated ribosomal protein gene expression—such as upregulation following inhibition of translation initiation and downregulation upon disruption of the Integrator complex or mRNA surveillance pathways—exhibited consistent effects with those observed in K562 and RPE cells.

Together, these results demonstrate that some core perturbation-response patterns are conserved across highly distinct cell types, suggesting the generalizability of these regulatory principles broadly across the cell-state landscape. In the meantime, we also observed distinct context-specific rewiring of some stress response pathways.

Because hESCs represent a transcriptionally and developmental very different cell state as compared to K562 and RPE cells, we also applied our few-shot perturbation prediction to hESCs to assess generalizability of this approach. The few-shot prediction performance on hESCs was again substantial improved as compared to the zero-shot model and the simple baseline models, starting from as few as ten shots (i.e. K = 10 input perturbations) and the accuracy increased as K grew (**Figure 6G**). As in the analyses for K562 and RPE cells, prediction accuracy for any fixed K value strongly depended on the magnitude of the perturbation effect, reflecting reduced signal-to-noise ratios for weaker perturbations (**Figure 6H**).These results suggest a generalizability of our few-shot prediction framework and supports the ability of the mixture-of-experts model to disentangle intrinsic gene functions from context-specific effects. Our comparison of the prediction accuracies between RPE1, K562 and hESC also suggest that perturbation prediction may be more difficult for cells with higher multipotency or pluripotency, potentially due to their stronger differentiation potential and more diverse differentiation outcome.

### Perturbation-induced differentiation of embryonic stem cells

Pseudo-bulk analyses revealed that multiple differentiation-associated programs were induced in the hESC CRISPRi screen (**Figure 6E**). Because differentiation often produces heterogeneous mixtures of cell states, single-cell resolution is required to resolve these transitions. To characterize perturbation-induced differentiation at the single-cell resolution, we leveraged our global reference cell-state manifold to determine which physiological states the perturbed cells most closely resembled.

We performed single-cell clustering using the Leiden algorithm and excluded clusters enriched for non-targeting sgRNAs to focus on genetically perturbed cells (**Figures 7A and Extended Data Figure 5A, B**). Clusters were annotated and grouped based on a combination of (i) the mean expression z-scores of the gene-expression programs in these clusters (**Extended Data Figure 6**), (ii) the enrichment of specific Gene Ontology^65^, Reactome^66^, CORUM^67^ and KEGG^68^ gene sets in the perturbed genes enriched in the clusters (**Supplementary Table 1**), and (iii) the zero-shot projection the cells in the clusters onto the global reference cell-state manifold (**Methods**). Most of these clusters correspond to stress-response states that preserved pluripotency and projected to epiblast-like states (**Figure 7B**). Notably, we identified three classes of clusters that strongly downregulated pluripotency while activating differentiation programs (**Figures 7A-C and Extended Data Figure 5C**): (1) clusters projecting to early embryonic cell types spanning all three germ layers (**Figure 7B,C**); (2) clusters projecting to mesenchymal cell types such as fibroblasts and smooth muscle cells (**Figure 7B,C**); and (3) clusters characterized by low UMI counts and high mitochondrial gene fractions, consistent with cell damage (**Extended Data Figure 5C**). Quantification of perturbation enrichments revealed 56 perturbed genes enriched in the mesenchymal differentiation class (grouping all mesenchymal differentiation clusters) and only 6 perturbed genes enriched in the germ-layer differentiation class (grouping all germ-layer differentiation clusters) (**Extended Data Figure 5D**).

**Figure 7.**
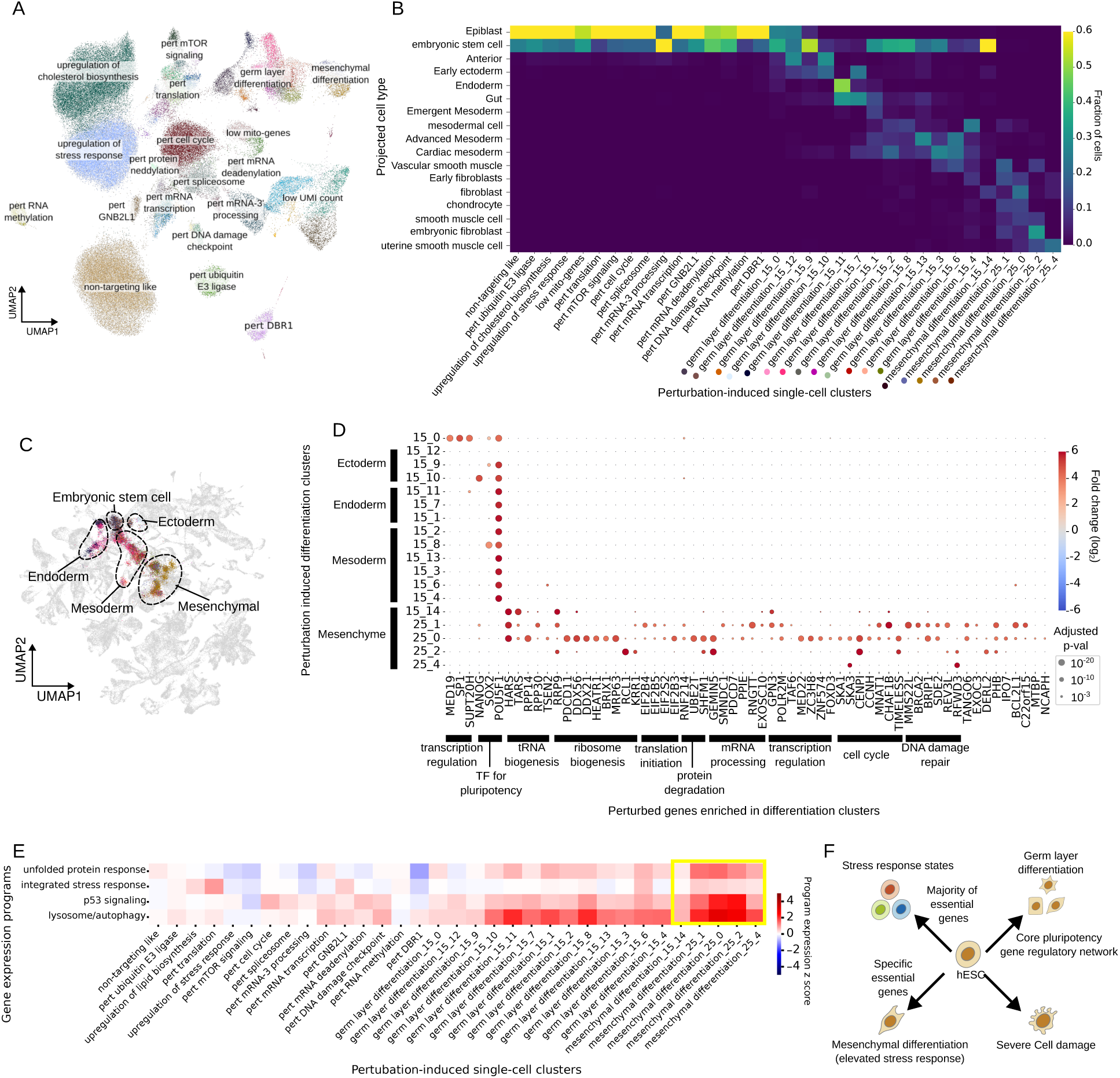
| Single-cell analysis of the genome-scale perturb-seq experiment in hESCs. **A,** UMAP of perturbed cells colored by Leiden cluster and labeled by the annotated classes. Only clusters enriched for gene-targeting sgRNAs are shown. Clusters are annotated by overrepresented perturbed-gene sets (“pert–…”) or perturbation-induced phenotypes. **B,** Confusion matrix comparing perturbation-induced single-cell clusters with projected cell types on the global cell-state manifold. **C,** Projection of differentiation-related clusters onto the global cell-state manifold. Reference cells in the global manifold are shown gray, and projected cells are colored by cluster identity. **D,** Enrichment of perturbed genes across differentiation-related clusters. Dot size encodes adjusted *p* value and color encodes enrichment ratio. Enriched perturbed genes (x-axis) are grouped by functional pathway (separated by thick horizontal lines). Differentiation-related clusters (y-axis) are grouped by major lineages (separated by thick vertical lines). **E,** The z-scores of the expression levels of four stress response gene programs for the single-cell clusters derived in (A). The yellow box highlights theclusters associated with mesenchymal transitions. **F,** Schematic summary of four different types of cell-state transitions induced by genetic perturbations identified in hESC experiment.

Within the differentiation-associated classes, we examined enrichment ratios and statistical significance for individual perturbations (**Figure 7D**; **Methods**). Perturbations of the core pluripotency factors POU5F1, SOX2, and NANOG were all enriched in the germ-layer differentiation class (**Figure 7D**). POU5F1 knockdown was broadly enriched across most clusters within this class, whereas SOX2 and NANOG perturbations were preferentially enriched in specific subpopulations—for example, NANOG loss favored ectoderm-like states, whereas SOX2 loss favored mesoderm-like states.

These patterns are consistent with models in which POU5F1, SOX2, and NANOG differentially bias lineage potential, and in which pluripotency emerges from a balance among these factors^93^. Notably, the observation that POU5F1 knockdown produced differentiation trajectories spanning all three germ layers contrasts with the prevailing view that POU5F1 primarily promotes mesodermal fates^93^, suggesting a broader role of POU5F1 in early lineage control. We also note that additional regulatory constraints present in vivo but absent in vitro may modulate these outcomes. Beyond the canonical pluripotency factors, perturbations of three transcriptional regulators—MED19 (Mediator complex), SUPT20H (SAGA complex), and SP1—were enriched in an “early-exit” differentiation cluster (**Figure 7D**). Given their broad roles in transcriptional regulation and chromatin modification, disruption of these factors may destabilize the pluripotency network and promote partial or premature differentiation.

In contrast to germ-layer differentiation, mesenchymal states were induced by perturbations spanning a wide range of essential cellular pathways, including DNA damage repair, transcription, mRNA processing, translation, and protein degradation (**Figure 7D**). This breadth suggests that the observed mesenchymal transitions arise as a generalized response to cellular stress rather than from disruption of specific lineage-specifying regulators. Consistent with this interpretation, these mesenchymal clusters exhibited particularly high expression of stress-response programs (**Figure 7E**).

Notably, among the eukaryotic initiation factors (eIFs), only EIF2B3, EIF2B4, EIF2B5, and EIF2S2—all components of the eIF2B–eIF2 complex^94^—were significantly enriched (supported by at least two sgRNAs each) among perturbations driving mesenchymal clusters, whereas other eIFs were instead enriched in translation-related stress clusters but not in mesenchymal differentiation clusters (**Figure 7D,Extended Data Figure 5E, F**). Because the eIF2 complex is a central regulator of ISR^79^, this specificity strengthens the link between ISR activation and the mesenchymal transition observed here. Supporting with this connection, ISR signaling has also been implicated in promoting mesenchymal transitions in cancer^95^.

Together, these observations suggest that cellular stress is a key molecular driver of perturbation-induced mesenchymal transitions in hESCs. This response pattern suggest that mesenchymal transitions may be a general “escape” mechanism^96^ that enables cells to adopt an alternative, stress-tolerant state under adverse conditions.

Our data and analysis thus revealed distinct classes of cell-state transitions in hESCs following genetic perturbations (**Figure 7F**). First, perturbations of essential genes predominantly elicit stress-responses with minimal effects on differentiation and many of these stress responses are conserved across distinct cell types. Second, perturbations of core pluripotency factors and selected transcriptional regulators drive differentiation toward specific germ-layer-like states, indicating that individual pluripotency factors bias lineage potential in distinct ways. Notably, loss of POU5F1 induces differentiation across all three germ layers. Third, perturbations of diverse essential cellular pathways trigger transitions into mesenchymal states, likely reflecting a stress-associated escape mechanism of the cell. Finally, a subset of perturbations caused overt cell damage and loss of pluripotency marker expression.

## DISCUSSION

In this work, we introduce a framework for delineating and understanding transcriptionally-defined cell-states and perturbation-induced cell-state changes at a global scale. By constructing global and interpretable representations of cell states and perturbations, our framework enables cross-dataset analyses and knowledge extraction from single-cell and functional genomics experiments. The framework comprises three major elements: (i) construction of a global representation of the cell-state landscape, (ii) identification of major axes of cell-state transitions, and (iii) identification of major classes of genetic perturbations. This framework places cell states, state transitions, and perturbations into a shared coordinate system that enables direct comparison across datasets from different contexts, providing the prospect of a unified view of how genetic perturbations move cells through a global cell-state space. We applied this framework to systematically analyze diverse perturb-seq datasets, revealing both conserved patterns and context-specific rewiring of perturbation-induced cell-state changes. Moreover, we performed a genome-scale perturb-seq experiment in hESCs, which validated and broadened these understandings and unraveled novel stress-induced mesenchymal transitions.

A central element of this framework is a global cell-state map, generated by our single-cell manifolder generator (SCMG), a contrastive learning method that combines the strong batch-removal capabilities with scalability. By learning a representation in which cells colocalize according to biological similarity rather than technical origin, SCMG enables diverse datasets to be interpreted relative to a common atlas of cell states. It allows zero-shot dataset integration and cell-state projection and annotation, facilitating analysis and interpretation of new single-cell datasets, especially for understanding the relationships between cell states. This capability becomes increasingly important as single-cell datasets continue to proliferate across laboratories, assays, and organisms. In its current implementation, our global cell-state manifold prioritizes biological variations conserved between human and mouse, which may attenuate inter-species differences as a trade-off. This limitation can be addressed by training the SCMG model using single-species data, especially for the generation of human-specific global cell state manifold as more human datasets become available in the future.

Utilizing this global reference cell-state manifold, in conjunction with available perturbation datasets, we identified a set of ∼50 gene-expression programs, providing a basis for describing major dimensions of cell-state variations that are shared across diverse contexts. These programs allow cell-state changes to be decomposed into movements along conserved biological dimensions, which in turn enables comparison of perturbations measured in different cell types and experimental settings in a common language, although the dimensionality reduction may obscure finer-grained phenomena that remain biologically important.

This framework also enables a global and context-shared view of perturbations themselves. By learning functional embeddings of perturbed genes shared across context, we move toward a data-driven organization of gene function based on transcriptional consequences observed in diverse datasets. Importantly, this representation explicitly accommodates context dependence while still capturing intrinsic functional similarity among genes. As additional perturbation datasets become available, such embeddings can provide a foundation for refining perturbation taxonomies and for systematically identifying which pathway behavior is conserved versus rewired across cell states.

One immediate implication of this representation is the feasibility of few-shot perturbation prediction. Rather than requiring dense perturbation data in every cell type of interest, our framework suggests that strategic, sparse sampling of perturbations across the functional space may be sufficient to enable prediction of perturbation effects of all genes in new contexts. This reframes the challenge of perturbation modeling from exhaustive data collection toward principled experimental design guided by coverage of cell-state and perturbation space.

Our application of this framework across Perturb-seq datasets from different cell types revealed several systems-level properties of cell-state regulation: cell differentiation programs are generally insulated from perturbations of essential cellular processes; stress-response pathways exhibit distinct stimulus spectra and pronounced context dependence: only a subset of essential processes is tightly coupled to proliferative capacity. In particular, p53 and lysosome/autophagy pathways are activated by nearly all perturbation classes, functioning as general stress responses, whereas UPR and ISR were selectively activated by specific perturbation types consistent with canonical UPR and ISR mechanisms ^79,84^. The ribosome-biogenesis gene program is co-regulated with cell proliferation, whereas ribosomal protein (RP) gene program follows a distinct regulation mechanism, which is selectively up- or down-regulated by different stimuli, significantly revising previous models ^86,88,89^. We also uncovered substantial cell-type-specific rewiring of stress responses.

We further tested these findings in a developmentally distinct context by performing a genome-scale perturb-seq experiment in hESCs and revealed new features of perturbation responses. Despite their high differentiation potential, hESCs largely preserve insulation between essential and differentiation programs. While hESCs and other cell types exhibited many shared pathway-level patterns of stress responses, hESCs display higher resistance to many perturbation-induced stresses, which echoes previous studies of individual stresses^90,91^ and broadens the scope of stress resistance of hESCs. In addition, hESCs exhibit heightened sensitivity to mitochondrial dysfunction. Notably, we observed a class of mesenchymal transitions in hESCs that emerges from diverse classes of genetic perturbations that affect core cell-stress pathways, suggesting that hESC engages mesenchymal transitions as a general stress escape mechanism. Epithelial mesenchymal transition (EMT) has been previously suggest as a mechanism for cancer cells to escape from an unfriendly environment^96^. Whether our findings of stress-induced mesenchymal transitions in hESCs are related to the escape mechanism found in cancer cells remains an open question.

Together, our findings highlight how conserved regulatory principles coexist with context-specific adaptations, and underscore the importance of studying perturbation responses across diverse datasets covering different regions of the cell-state landscape. Moving forward, this work points toward a principled approach to perturbation predictions. Currently, the accuracy of generalizable perturbation prediction models is limited by the low coverage of starting cell states across the cell state space. Our global representations of cell states and perturbations provide a substrate onto which new datasets can be continuously mapped and interpreted, enabling systematic assessment of what is conserved, what is context specific, and where biological knowledge remains sparse. Because perturbation effects are strongly context dependent, data must be acquired across diverse cell states. Our global cell-state manifold also offers a natural guide for experimental design: perturbation screens can be prioritized in representative cell states spanning major identity domains, followed by denser sampling of the transitional regions where regulatory behavior changes most rapidly. In addition, our functional embedding of genes and few-shot prediction approach also allow principled and efficient design of perturbations, strategically choosing perturbed genes to span major domains in the functional embedding space without the need of dense sampling. Model performance can then serve as a quantitative criterion for data sufficiency. As predictive accuracy improves, this strategy can be extended beyond single-gene perturbations to higher-order combinatorial genetic perturbations^97^ and environmental changes by small molecules^98^, engineered regulators^99^, and physiological condition changes. Global perturbation modeling is not a static endpoint but an iterative process^100,101^, in which data collection and model refinement jointly expand the predictive scope of perturbation biology, and the framework reported in this work can provide a foundation to accelerate this process.

## Data availability

The down-sampled representation dataset of the global cell-state manifold and the standardized pseudo-bulk perturbation dataset are available at https://huggingface.co/datasets/xingjiepan/SCMG_data/tree/main/data. The global gene expression patterns can be visualized with a web browser at https://xingjiepan.github.io/SCMG. The genome-scale hESC perturb-seq data is available at NCBI GEO data repository (accession number: GSE295214).

## Code availability

The code for the SCMG package is available at https://github.com/xingjiepan/SCMG. The analysis scripts for generating figures in this paper is available at https://github.com/xingjiepan/SCMG_scripts/tree/main/manuscript2025.

## Acknowledgements

We thank Cheen Euong Ang, Karina Smolyar, and all members of the Zhuang and Weissman labs for the discussions. X.P. is a Howard Hughes Medical Institute Jane Coffin Childs postdoctoral fellow. R.A.S. is a Harvard Junior Fellow. J.S.W and X.Z. are Howard Hughes Medical Institute investigators.

## Author contributions

X.P., R.A.S., J.S.W. and X.Z. conceived the project. X.P. developed the computational framework. R.A.S. and J.M.R. designed and performed the hESC genome-scale perturb-seq experiment. X.P. and R.A.S. processed the perturb-seq data. X.P. performed the data analysis. X.P. and X.Z wrote the paper with input from R.A.S. and J.S.W..

## Additional information

**Supplementary information** is linked to the online version of the paper.

**Correspondence and requests for materials** should be addressed to Xiaowei Zhuang and Jonathan Weissman

## Supplementary Figures

**Extended Data Figure 1.**
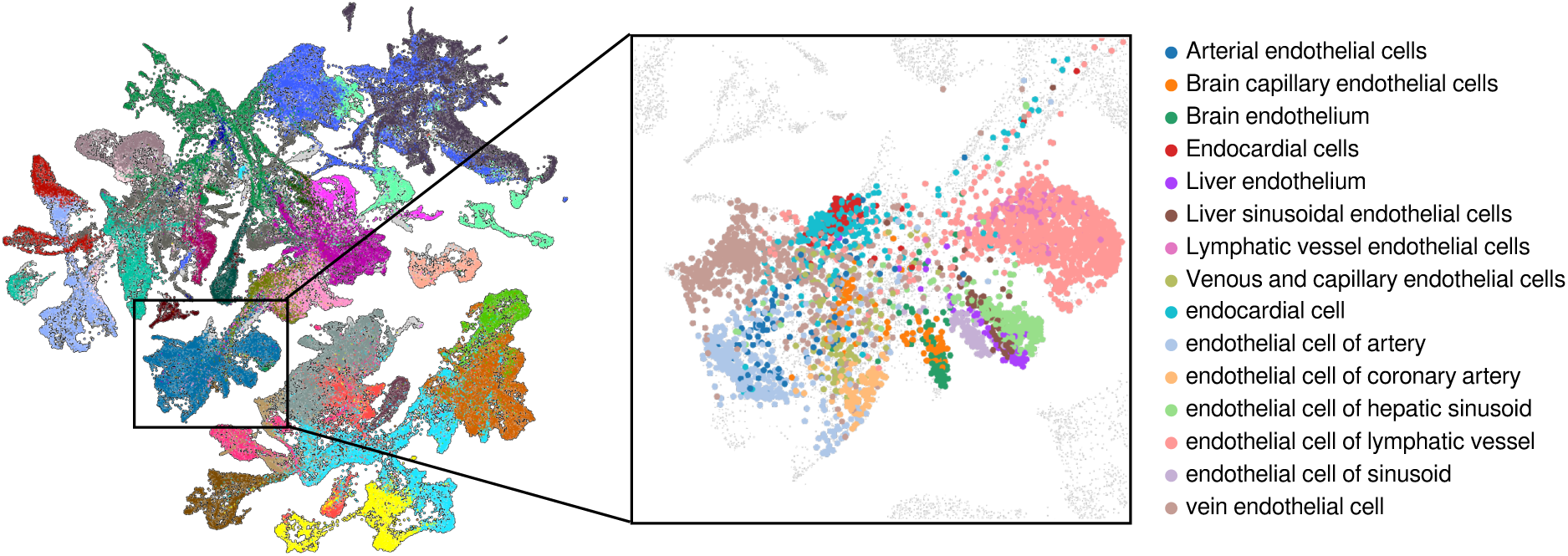
| Global cell-state manifold at sub-cell-type resolution, using endothelial cells as an example. Global UMAP of the cell-state manifold (left) colored by major cell types, with a zoomed-in view of the endothelial region (right). In the inset, exemplar endothelial subtypes are highlighted by colors, and all other cells are shown in gray.

**Extended Data Figure 2.**
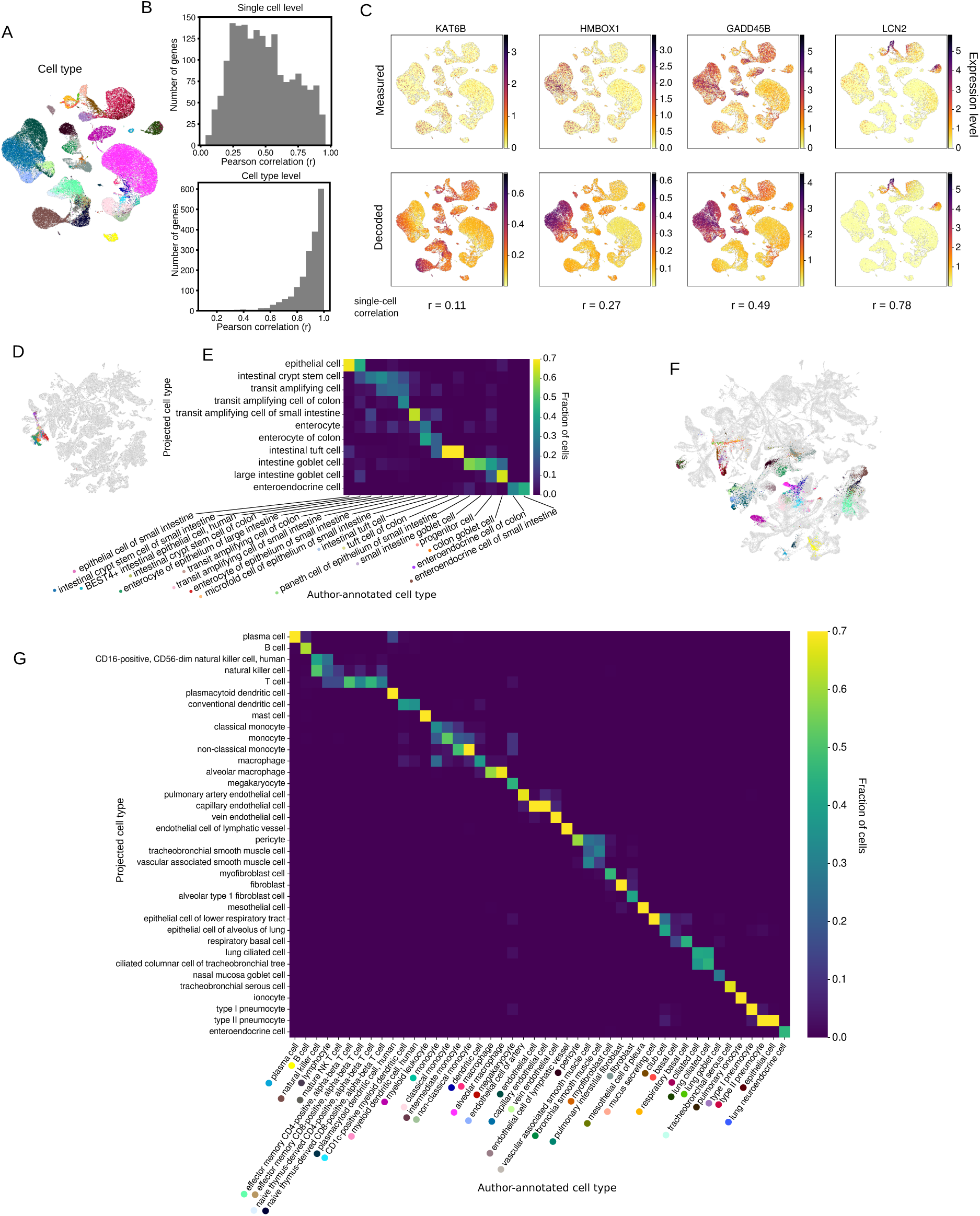
| Decoding and zero-shot projection by SCMG. **A,** UMAP of lung cells from Ref. ^55^, a dataset unseen during SCMG training, colored by author-annotated cell types. **B,** Distributions of Pearson correlation coefficients between originally measured and SCMG-decoded expression values for individual genes across cells (top) and across cell types (bottom). **C,** Examples of measured (top) and decoded (bottom) expression for four representative genes visualized on UMAP. Correlation coefficients between measured and SCMG-decoded expression levels across cells are indicated below the decoded panels. **D,** Zero-shot projection of the intestine dataset from Ref. ^54^ onto the global cell-state manifold. Reference cells in the global manifold are shown in gray. Projected cells are colored by their author-annotated types as in (**E**). **E,** Confusion matrix comparing author-annotated intestine cell types with SCMG-projected types. Only the top projected type from projection for each author-annotated cell type is shown. **F,** Zero-shot projection of lung cells from Ref. ^55^ onto the global cell-state manifold. Reference cells in the global manifold are shown in gray. Projected lung cells are colored by author-annotated types as in (**G**). **G,** Confusion matrix comparing author-annotated lung cell types with SCMG-projected types. Only the top cell type from projection for each author-annotated cell type is displayed.

**Extended Data Figure 3.**
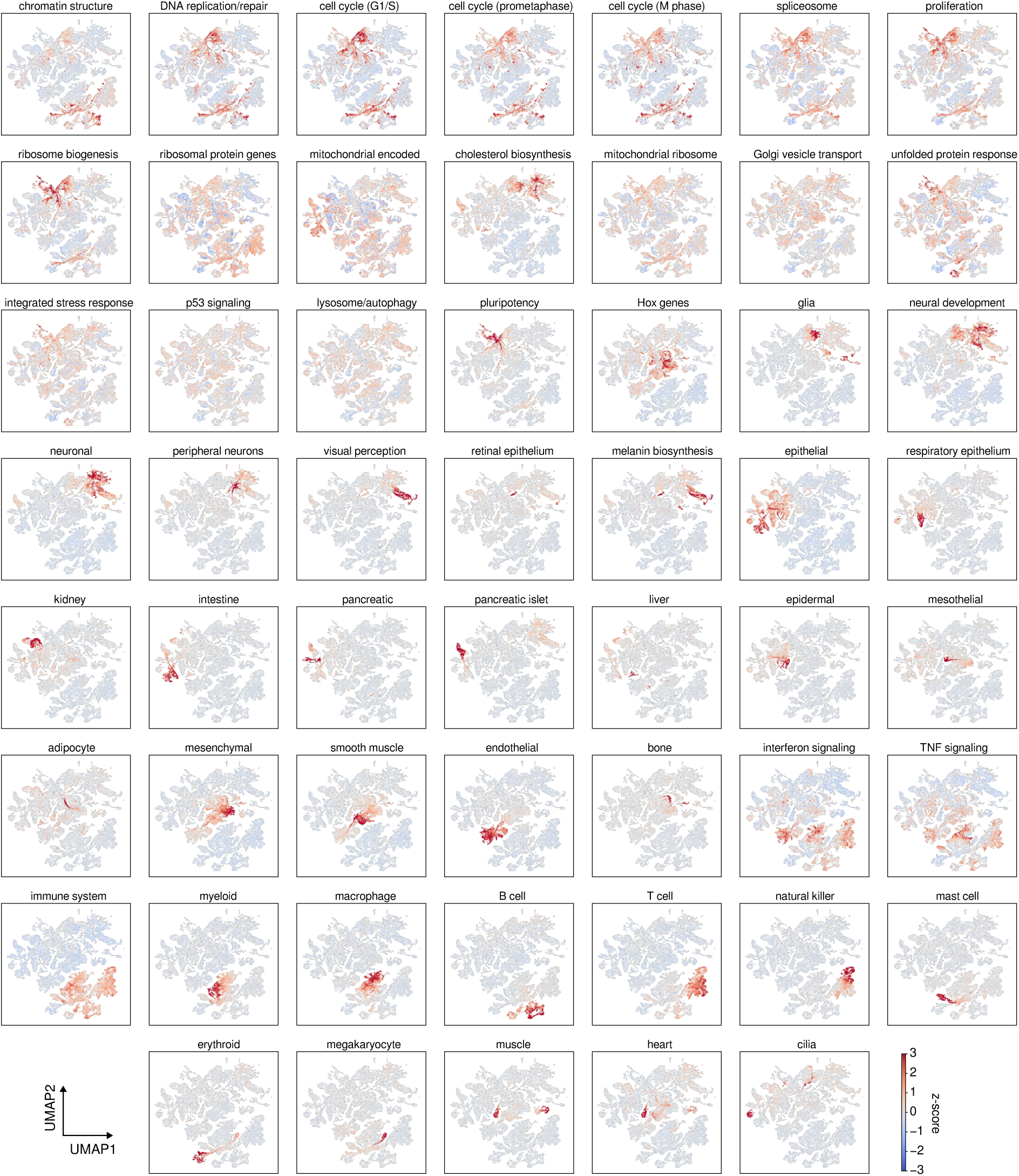
| Mean expression of the 54 gene-expression programs across the global cell-state manifold. Each UMAP plot shows the expression level of individual gene-expression programs (defined as the average expression of its member genes) across the global cell-state manifold.

**Extended Data Figure 4.**
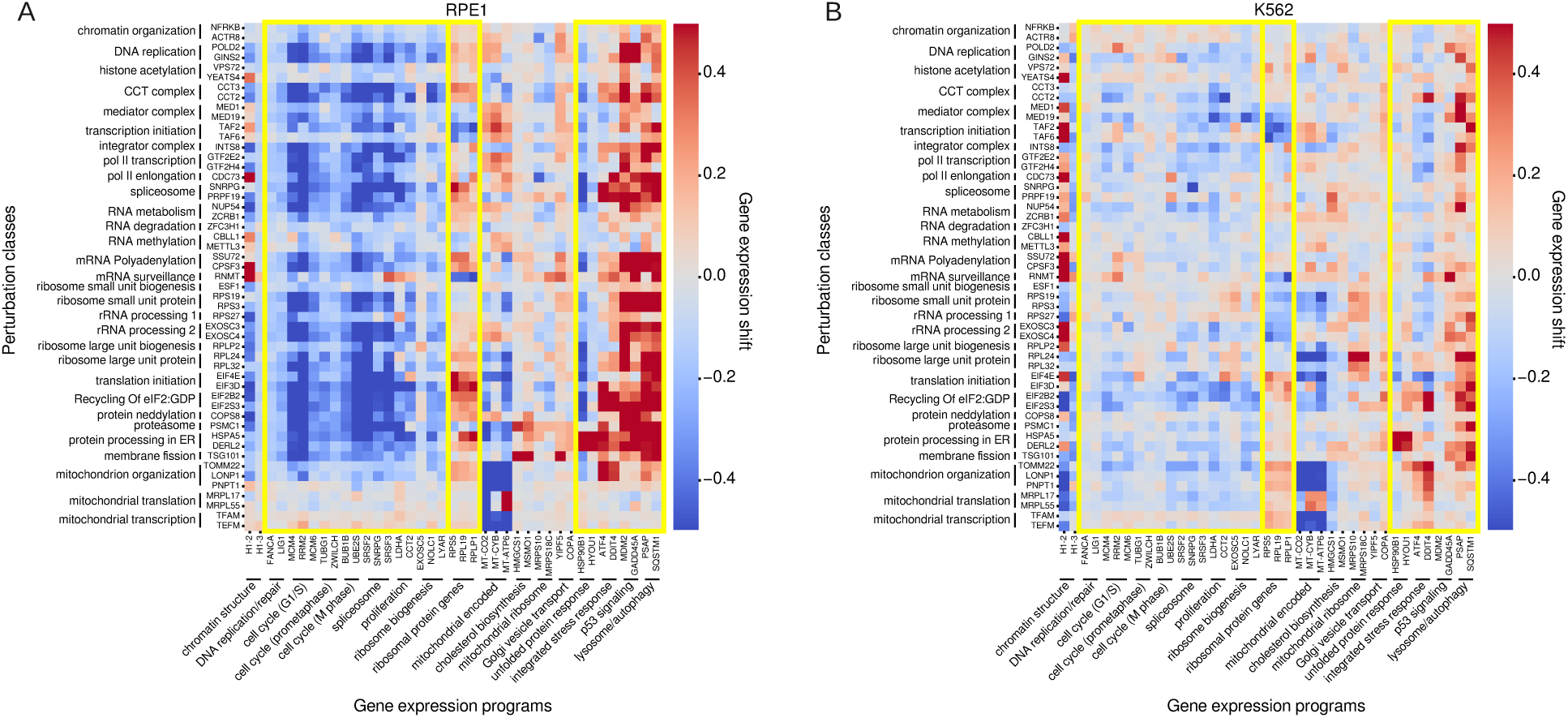
| Expression changes of marker genes of essential programs induced by perturbing marker genes in the perturbation classes. One to three marker genes were selected for each gene expression program (marked by thick horizontal lines) or perturbation class (marked by thick vertical lines). Yellow boxes highlight gene expression programs related to cell proliferation, ribosome protein genes, and stress response. **A,** RPE1 CRISPRi screen. **B,** K562 CRISPRi screen.

**Extended Data Figure 5.**
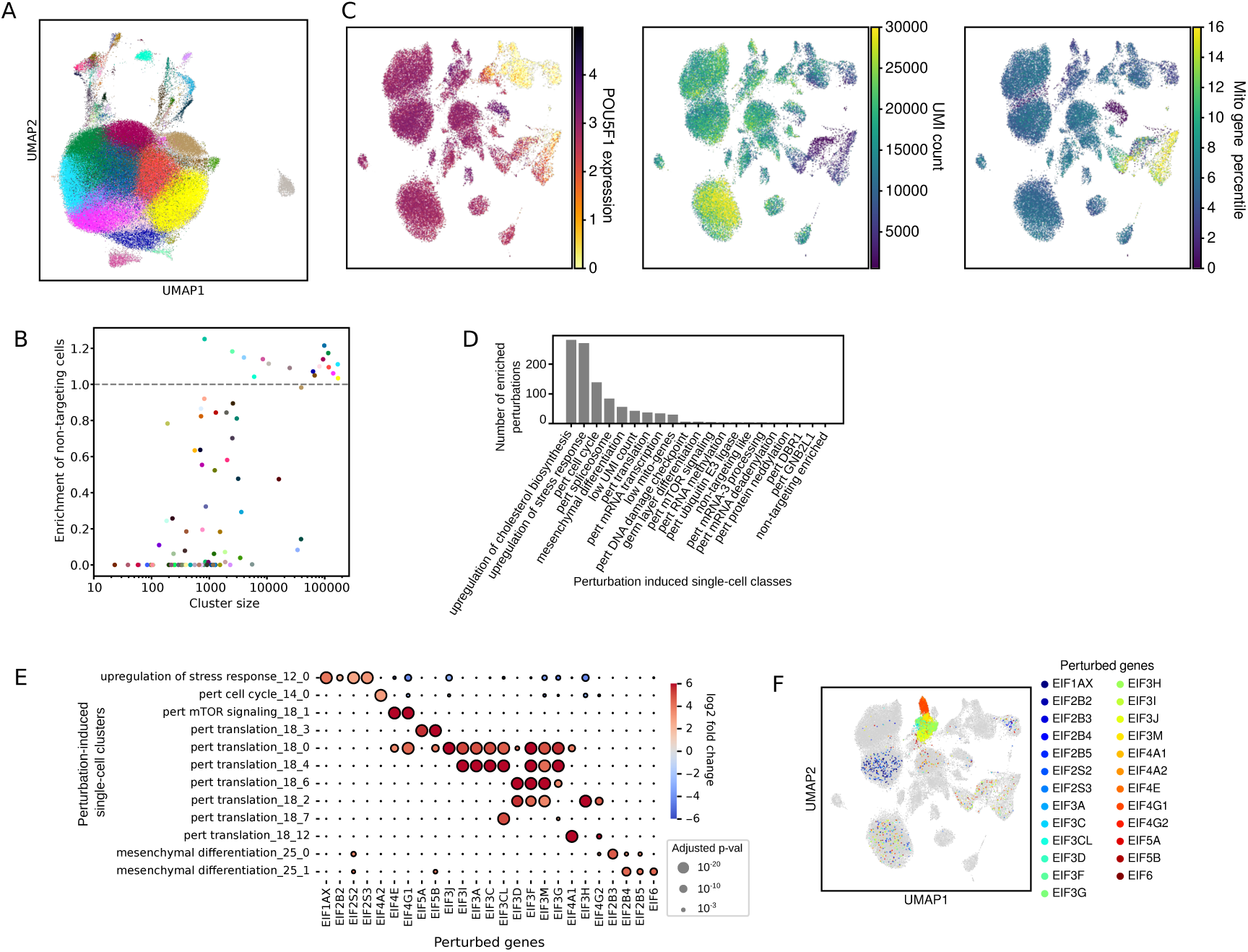
| Additional single-cell analyses of the genome-scale hESC Perturb-seq dataset. **A,** UMAP of all cells from the screen, colored by Leiden cluster. **B,** Scatter plot showing, for each cluster, its enrichment ratio of cells carrying non-targeting sgRNAs versus its size (number of cells). **C,** UMAPs displaying POU5F1 expression (left), per-cell UMI counts (middle), and mitochondrial-gene percentage (right) of the perturbed cells. **D,** Number of perturbed genes significantly enriched in each perturbation-induced single-cell class (groups of related clusters). **E,** Perturbation-induced single-cell clusters enriched for perturbations targeting eIF genes. Dot size represents adjusted p-values; color indicates enrichment ratio. We note that although EIF5B and EIF6 showed enrichment in mesenchymal differentiation clusters, their enrichment was only observed for a single sgRNA, unlike EIF2B3, EIF2B4, EIF2B5, and EIF2S2, which were all evidenced by at least two sgRNAs. **F**, UMAP of perturbed cells. Cells targeted by eIF gene perturbations are colored by the specific perturbed gene; all other cells are shown in gray.

**Extended Data Figure 6.**
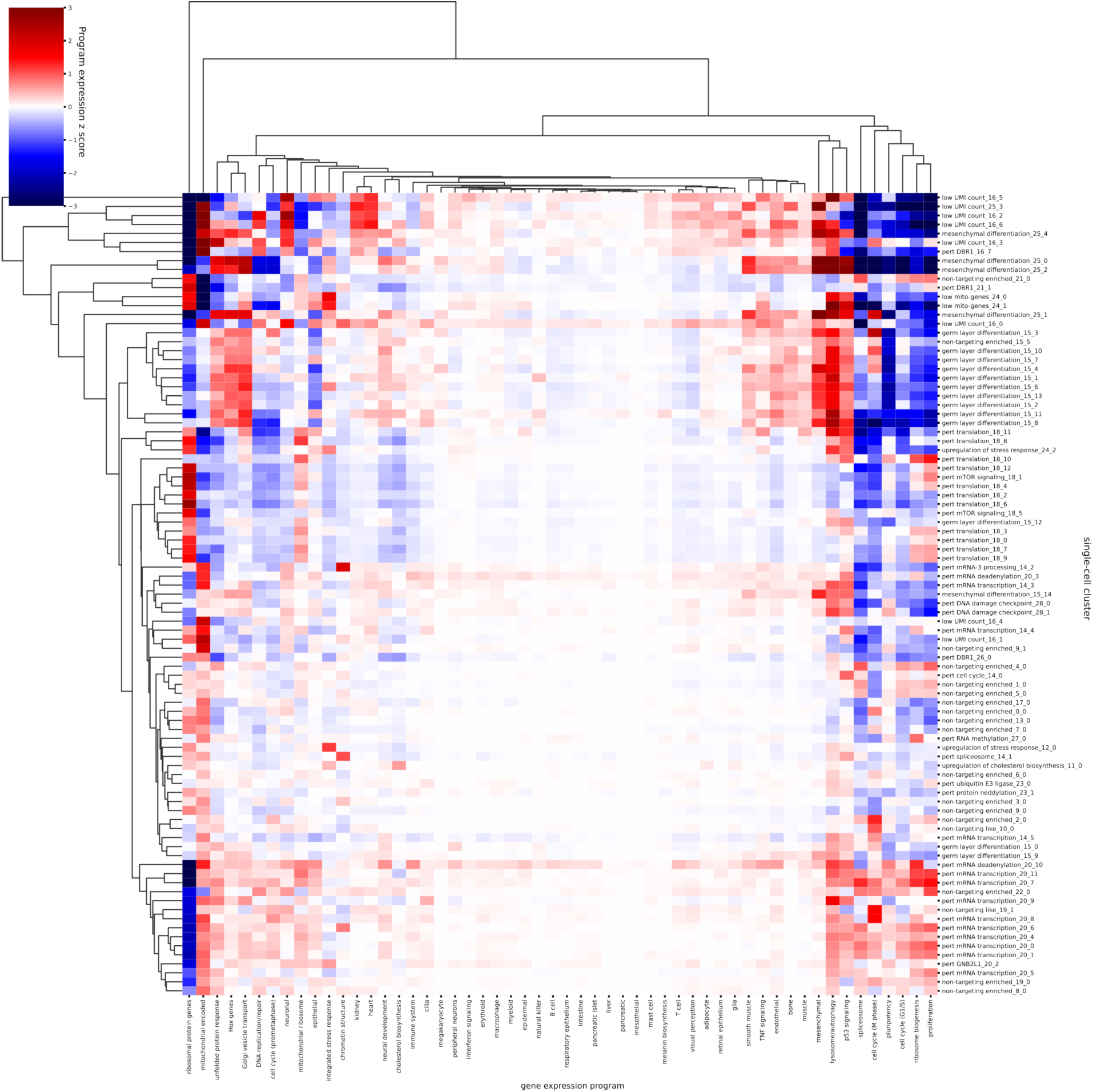
| Shift scores of gene-expression programs for individual single-cell clusters in the hESC Perturb-seq experiment. Mean gene-expression program shift scores for each perturbation-induced single-cell cluster are shown. The shift scores are normalized by the average score of the clusters enriched for non-targeting guides.

## Supplementary table captions

**Supplementary Table 1 | Significantly enriched perturbed genes in the perturbation-derived single-cell classes of the hESC perturb-seq experiment.** Each row is a pair of a single-cell class and a perturbed gene. The “class” column is the name of the single-cell class. The “enriched_perturbed_gene” column is the perturbed gene enriched in the class. The “n_enrcihed_guides” column is the number of guide RNAs supporting the enrichment.

**Supplementary Table 2 | The scRNA-seq datasets for training and benchmarking the SCMG model.** Each row is a dataset. The “dataset_id” column is the dataset identifier. The “species” column is the species measured in the dataset. The “used for SCMG training” column is whether the dataset was used for training the SCMG model. The “reference dataset” column is whether the dataset was used as a reference for pairwise integration. We selected 5 large datasets with diverse cell types as the reference datasets and integrated other datasets to the reference datasets. The “cell count (after subsample)” column is the number of cells in the dataset after subsampling that are used for training the SCMG. The “reference” column is the reference number with which the dataset is cited in the manuscript.

**Supplementary Table 3 | The annotated pairs of cell type labels from different datasets corresponding to similar cell types.** Each row is a pair of similar cell types. The “dataset1” column is the dataset containing the first cell type. The “dataset2” column is the dataset of containing second cell type. The “cell_type1” column is the name of the first cell type. The “cell_type2” column is the name of the second cell type.

**Supplementary Table 4 | The annotated major cell type labels of cell types.** We manually grouped the author-annotated cell types into 31 major cell types. Each row is a cell type. The “cell_type” column is the cell type label. The “major_cell_type” column is the major cell type label.

**Supplementary Table 5 | The collection of datasets of RNA-sequencing before and after genetic perturbations.** Each row is a dataset. The “dataset” column is the dataset name. The “species” column is the species measured in the dataset. The “type” column is the experiment type (single-cell or bulk measurements). The “number of perturbations passed QC” column is the number of perturbation entries in the dataset that passed the quality control. The “reference” column is the reference number with which the dataset is cited in the manuscript.

**Supplementary Table 6 | The oligonucleotide pool for the guide RNAs used in the hESC perturb-seq experiment.** Each row is a synthesized oligonucleotide. The “element” column is the name of the oligonucleotide. The “sequence” column is the sequence of the oligonucleotide.

## Methods

### Single-cell manifold generator (SCMG)

The main goal of SCMG is to learn a generic strategy to map cells characterized by their transcriptome profiles (e.g. as measured by scRNA-seq) from diverse datasets to a universal representation space that accurately describes the cell states. Therefore, SCMG needs to keep biological variations while removing batch effects that cause cells of the same biological state to have different measured transcriptional profiles in different experiments. To achieve this goal, we designed the SCMG model as an autoencoder trained with contrastive learning. The encoder is a multilayer perceptron (MLP) neural network that maps a single-cell transcriptional profile vector *x_in_* to a 512-dimensional latent vector *z* = *encoder*(*x_in_*). Each point in the latent space represents a batch-corrected cell state. To make the encoder generic and work for any unseen datasets, we used the transcriptional profiles without any information about dataset identities as the inputs of the encoder. The decoder is an MLP that maps a cell-state vector *z* in the latent space back to a transcriptional profile. Since the same cell state can have different measured transcriptional profiles in different datasets, we also took the dataset identity *A* as an additional input for the decoder neural network to specify the dataset. Hence, the decoded transcriptional profile vector is a function of both the cell-state vector *z* and the dataset identity *A*: *x_decode_* = *decoder*(*z*, *A*). A mean squared error loss term *MSE*(*x_in_*, *x_decode_*) is used to minimize the information loss during the encoding-decoding process, so that the latent space can retain maximum biological variation.

Next, we aimed to minimize batch effects using a contrastive learning approach. In this approach, we first perform pairwise integration of datasets that describe similar or overlap cell-state distributions (see the “Pairwise integration of scRNA-seq datasets” section below). For each example pair of scRNA-seq datasets A and B, we identified inter-dataset pairs of cells that represent biologically similar cell states by co-embedding the pair of datasets (see the “Pairwise integration of scRNA-seq datasets” section) and identifying the mutual K nearest neighbors (mKNN, K = 10 in this study) pairs of cells between the datasets^102^. The mKNN pairs can reasonably approximate cells of similar biological states and we connected each of these cell pairs with an edge. We then considered a cell *x_i_* from the dataset A and a cell *x*_’_ from the dataset B that are biologically similar (i.e. connected with an edge), as well as a cell *x_j_* from the dataset B that is biologically different (unconnected) from *x_i_* and *x_j_*. Let the Euclidean distances among the three cells in the embedding space be *d_i,j_* = *dist*(*z_i_*, *z_j_*), *d_i,k_* = *dist*(*z_i_*, *z_k_*), and *d_j,k_* = *dist*(*z_j_*, *z_k_*). Our strategy for batch-effect removal is to achieve *d_i,j_* < *d_i,k_* and *d_i,j_* < *d_j,k_* for such cell triplets. To minimize the batch effects by training the model, we define a contrastive loss term as *L_contrastive_*(*i,j,k*) = *softplus*(*d_i,j_* − *d_i,k_*) + *softplus*(*d_i,j_* − *d_j,k_*). The training of SCMG should optimize the model’s latent embedding to minimize the total of such contrastive loss terms for all triplets of cells from all pairs of datasets.

The training data of SCMG contain the transcriptome-wide gene-expression vectors of all cells from the training datasets and pre-defined tuples (*i*, *j*, *A*, *B*) of cell pairs of similar biological states, where *i*, *j* are cell IDs and *A*, *B* are the respective dataset IDs. The training process iterates through the cell pair tuples. For each cell pair, we randomly sample a cell *k* from the dataset B to calculate the contrastive loss *L_contrastive_*(*i*, *j*, *k*). The minimization of the contrastive loss pulls cells *i*, *j* to be close to each other and pushes the cell *k* to be far apart. To regularize the magnitudes of the embedding vectors, we add the squared L2 norms of the embedding vectors to the loss function as 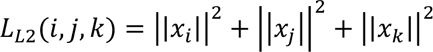. Then we compute the mean squared error loss for the cell *i* as *MSE*(*x_i_*, *x_i,decode_*). The total loss is defined as

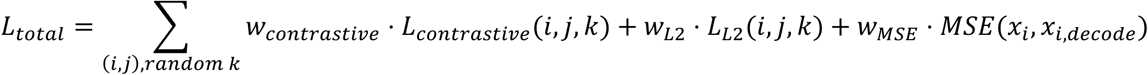

where *w_contrastive_*, *w*_*L*2_ and *w_MSE_* are the weights of the respective loss terms. To speed up the training, our sampling of the cell *k* was random, which means that the cell *k* may or may not be biologically similar to *i* and *j*. We note that randomly sampling *k* should give very similar training results as requiring *k* to be dissimilar from *i* and *j* when the model converges, because randomly picking a biologically different *k* is much more probable than picking a *k* of the same cell state as *i* due to the high diversity of cell states.

### Collection and preprocessing of scRNA-seq datasets

We collected 39 scRNA-seq datasets covering diverse cell types for humans and mice (**Supplementary Table 2**). We selected 36 datasets^13–46^ for training the SCMG model and left out 3 datasets^54,55,103^ to perform zero-shot integration benchmarking of the model’s performance. We down-sampled some of the datasets with large numbers of cells to avoid over-representation of certain cell types caused by the biased sampling. For mouse datasets, we mapped their gene IDs to their human ortholog gene IDs according to annotations from the Ensembl database^104^.

### Pairwise integration of scRNA-seq datasets

We performed pairwise integration to identify cell pairs measured in different datasets that comprise similar biological (gene expression) states. For each pair of datasets to be integrated, we used the cell types annotated by the dataset authors to identify pairs of cell types that represent biologically similar states. Because different studies often annotated cell types with different conventions, we manually inspected the annotations and identified matched cell-type pairs representing similar cell states. Due to the difference in the granularity of cell-type annotations, a cell type in one dataset could be matched to multiple cell types in another dataset. We removed the cells whose cell-type identity cannot be matched to the other dataset for pairwise integration purpose. For the remaining cells, we subset the genes to the common genes measured in both datasets. We normalized the total count of each cell to 10,000 and performed a log1p transformation of the cell-by-gene matrices. We then identified the top 2,000 highly variable genes of the combined dataset using the highly_variable_genes function from Scanpy^105^. We converted the normalized gene expressions to z-scores and used the highly variable genes to calculate a 100-dimensional PCA embedding vector for each cell. Using the PCA embedding, we found, for each cell in one dataset, the 10-nearest neighbors from the other dataset. Then we identified pairs of cells that were mutual neighbors. The cell-type labels for each mutual neighbor cell pair may or may not belong to the pre-defined cell-type pairs representing biologically similar states. We further filtered the mutual neighbor cell pairs to keep only those cell pairs whose cell-type labels belong to predefined cell-type pairs representing similar cell states. We defined the resulting cell pairs to have similar biological states and used these cell pairs to train the SCMG model.

The time cost for manually identifying the corresponding cell types between all pairs of datasets scales quadratically with the total number of datasets, making it impractical to manually annotate all possible dataset pairs from the 36 datasets. We thus selected 5 large datasets with diverse cell types as the reference datasets (**Supplementary Table 2**) and integrated the other datasets to these reference datasets, further requiring that the integrated dataset pairs contain at least some matching cell types. In total, we manually annotated 125 pairs of datasets encompassing 4,918 matched cell-type pairs (**Supplementary Table 3**). Integration of the dataset pairs resulted in 4,917,573 cross-dataset pairs of cells that belong to similar biological states. We also note that during integration, only cells with matching cell-type labels in both datasets in the pair were integrated.

We noticed some datasets had internal batch effects between different experiments within the same dataset. We thus performed intra-dataset integration of different batches and also included the cell pairs belonging to similar biological states identified between different experiments within the same dataset for SCMG training. We calculated intra-dataset cell pairs by identifying the 8 nearest neighbor cells in the PCA space. To emphasize the inter-dataset integration, we downsampled the datasets to 10% of cells when computing intra-dataset cell pairs. In total, the training dataset has 5,585,245 pairs of cells, including both inter-dataset and intra-dataset cell pairs.

### Define a standard gene set for human and mouse

First, we downloaded the table of all human genes and their mouse orthologous genes from the Ensembl database^104^ and selected the genes with ortholog annotations. Then, we filtered out the genes that did not have a symbol assigned by the Human Genome Organisation Gene Nomenclature Committee^106^. Finally, we filtered out the genes that were not measured in the Tabula Sapiens dataset, which is a reference scRNA-seq dataset that covered most human organs. The resulting standard gene set contained 18,108 genes. We used the standard gene set to integrate datasets from different studies.

### Training of the SCMG model

We merged all cell-by-gene matrices from the 36 training datasets into a matrix and kept only genes in the standard gene set. If a gene in the standard gene set is not measured in a training dataset, the expression values of the missing gene were set to 0 for cells from the dataset. We tracked the missing genes for each cell so that the missing genes were excluded from the calculation of the reconstruction loss term. We normalized the total expression count of each cell to 10,000 and performed a log1p transformation. For each epoch of training, we iterate through the pairs of biologically similar cells identified by pairwise integration. For each pair of cells, we calculated the loss function defined in the “Single-cell manifold generator (SCMG)” section. We optimized the model’s parameters by minimizing the loss function with stochastic gradient descent using the AdamW optimizer^107^. The training was performed for 100 epochs.

### Selection and annotation of the representation dataset

To visualize the cell-state organization in the global single-cell manifold, we subsampled cells from the training dataset to create a representation dataset with balanced numbers of cells from each cell type. For each dataset-author-annotated cell type from each dataset, we randomly selected 100 cells. If the cell type did not have more than 100 cells, all cells were kept. We merged the selected cells into a dataset and kept only genes the standard gene set. The expression levels of the unmeasured standard genes were set to 0. The representation dataset contained 133,061 cells of 797 cell type labels. We manually grouped the author-annotated cell types into 31 major cell types (**Supplementary Table 4**). We embedded the cells with SCMG and displayed the cells in a 2D UMAP using the SCMG embedding.

### Benchmarking of integration performance

We compared the performance of SCMG and other 5 deep learning models (scVI, Geneformer, scGPT, UCE, and SCimilarity) on zero-shot integration performance of scRNA-seq datasets using their respective cell-state embeddings. The embeddings of scVI, Geneformer, scGPT, and UCE were downloaded from the CELLxGENE database^108^ with the census version 2023-12-15. We calculated the SCimilarity embedding with its model v1.1^52^. For each pair of datasets to be integrated, we selected cell types that were measured in both datasets and co-embedded the selected cells with each deep learning method. Then we computed a 30-nearest-neighbor graph for the co-embedded cells (each of these nearest-neighbor cell pairs are connected by an edge) and visualized the co-embedded manifolds by plotting a 2D UMAP. We quantified the integration performance using this nearest-neighbor graph. For each edge in the nearest-neighbor graph, we compared the dataset labels of the two cells connected by the edge. If the two cells were from different datasets, we classified the edge to be an inter-dataset edge. We calculated the fraction of inter-dataset edges among all edges to quantify the performance of batch effect removal. Next, we quantified the preservation of biological variation using the cell type labels. Because the same cell state can have different labels in different datasets due to different annotation convention, we only consider intra-dataset edges for this analysis. We compared the cell-type labels of the two cells connected by each edge and classified the edge to be a same-cell-type edge if the two cells have the same cell type labels. We quantified the preservation of biological variation by calculating the fraction of same-cell-type edges among all edges. For each of these two metrics, to calculate the significance of performance difference between different deep learning methods, we calculated the metric values for all pairs of benchmarking integration datasets and computed the p-value using two-sided Welch’s t-test.

### Benchmarking of decoding performance

For each benchmarking dataset, we identified highly variable genes using the highly_variable_genes function from Scanpy^105^. We calculated the decoded gene expressions for each cell to get a decoded cell-by-gene matrix. Then we filtered both the measured cell-by-gene matrix and the decoded cell-by-gene matrix to only include the highly variable genes. For each gene, we calculated the Pearson correlation coefficient between the vector of measured expressions of this gene across individual cells and the corresponding vector of decoded expressions. The Pearson correlation coefficient quantifies the decoding accuracy at single-cell level. Then we calculated the mean expression level for each gene in each cell type to get cell-type-by-gene matrices for the measured and decoded gene expressions. We then calculated, for each gene, the cell-type level Pearson correlation coefficients between measured and decoded gene expressions across different cell types.

### Projection of query cells onto the representation dataset

Given a query cell defined by its transcriptional profile, we removed the genes that are not in the standard gene set and set the expression level of the missing standard genes to zero. We normalized the total count to 10,000, log1p transformed the expression vector and used the SCMG model to calculate the 512-dimensional cell-state embedding. We identified the closest cell from the representation dataset by Euclidean distance and defined the closest cell as the projected cell state of the query cell. To visualize the query cell on the global cell state UMAP, we assigned the 2D UMAP coordinate of the projected cell state plus a small Gaussian random shift to the query cell. The addition of a random shift was to visually separate multiple cells that were projected onto the same cell state, so that cell densities could be better visualized by the cloud of points.

### Collection and preprocessing of published perturbation datasets

We collect 9 datasets of RNA-sequencing before and after genetic perturbations (**Supplementary Table 5**). The datasets were downloaded from their respective sources as specified in their paper or from the scPerturb database^109^. We filtered the single-cell datasets with pooled perturbations for high quality perturbations as follows. We only kept the perturbations that had at least 50 cells. For perturbations that aimed to knockdown gene expressions, we compared the gene-expression levels in the perturbed cells versus the control cells and selected the perturbations whose target genes’ mean expression levels in the perturbed cells were <80% of the mean expression levels in the control cell. We also required the perturbed gene expression levels to be significantly different between the perturbed cells and the control cells using the two-sided Mann–Whitney U test with a p-value threshold 0.01. For perturbations that overexpress genes, we filtered the perturbations with a similar strategy by requiring the fold change of gene expression to be greater than 1.5 and the p-value less than 0.01. We aggregated the single-cell gene expressions into pseudo-bulk expressions by calculating the mean expression values for cells with the same perturbations for each dataset.

For each dataset, we kept only genes in the standard gene set (defined in the “Define a standard gene set for human and mouse” section) and set the expression levels of missing standard genes to 0. For each bulk or pseudo-bulk expression profile, we normalized the total count to 10,000 and performed a log1p transformation. We defined the perturbation-induced gene expression shift as the difference between the normalized expression vector for the perturbed state and the normalized expression vector for the control state. We combined the standardized bulk/pseudo-bulk perturbation-induced gene expression shift vectors as a reference database of perturbation effects.

### Compute human/10x-equivalent decoded single-cell transcriptomes

To obtain a human/10-equivalent decoded expression profile for any cell, we first encoded the cell’s transcriptome with the SCMG encoder to generate its latent embedding. Second, within a global reference compendium, we identified the cell’s nearest neighbor among only human cells profiled by the 10x assay and recorded that neighbor’s dataset ID. Finally, we decoded the query cell’s latent embedding with the SCMG decoder while conditioning on the dataset ID of this nearest-neighbor human/10x cell, yielding the human/10-equivalent decoded gene-expression vector. This procedure removes species- and assay-specific effects by mapping all cells to their human/10x equivalents.

### Compute gene expression entropy

Using human/10-equivalent decoded gene-expression vector from the reference compendium (containing the 36 scRNA-seq datasets), we computed an expression entropy score for each gene. For a given gene *g*, we first calculated its mean expression within each author-annotated cell-type cluster, yielding a cell-type-level vector *x_g_*. We then L1-normalized this vector to obtain a discrete, distribution-like vector *p_g_* = *x_g_* / ∑_*i*_*x_g,i_*, where *i* is the indices representing cell types. The entropy of the gene’s expression profile was defined as *H_g_* = − ∑_*i*_ *p_g,i_* log (*p_g,i_*), with higher values indicating more uniform expression across cell types.

### Global identification of gene expression programs

We identified programs by jointly analyzing batch-corrected (human/10x equivalent) global profiles and perturbation-induced pseudobulk shifts. The batch-corrected and pseudobulk shift values represent gene expression values at the log1p scale (see “Training of the SCMG model” and “Collection and preprocessing of published perturbation datasets” sections). As a first step, we filtered genes using their global expression profiles. We removed genes whose peak mean expression (i.e. expression level in their highest-expressing cell type) was <0.2, and excluded genes with expression-entropy >6.4 to retain those differentially expressed across cell types. For each remaining gene, we defined its expression vector as the gene’s z-score normalized expression levels across all single cells in the reference compendium. We computed pairwise Pearson correlations between the gene expression vectors and, for each gene, identified “highly correlated partners” with r > 0.5. Genes with fewer than five partners were discarded, as they did not form coherent co-expression clusters with other genes. The expression vectors of the retained genes were then reduced to a 50-dimensional space using principal component analysis (PCA).

For the perturbation data, we selected a set of readout genes (as described below) and defined, for each readout gene, a perturbation response vector as its pseudobulk expression shifts across 11,406 perturbation conditions. We selected readout genes showing strong responses—absolute pseudobulk shift ≥ 0.5 in at least five perturbation conditions—then computed pairwise Pearson correlations between response vectors. For each gene, highly correlated partners were defined as those with r > 0.5; genes with fewer than five partners were removed. Response vectors for the remaining readout genes were reduced to a 50-dimensional space using PCA.

To weight physiological and perturbation signals equally, we scaled the perturbation PCA embeddings so that the element-wise standard deviation matched that of the physiological PCA embeddings. For each gene, we then concatenated its scaled perturbation PCA vector with its physiological PCA vector. If a gene met filters in only one modality, the missing modality’s PCA vector was set to zero. In this 100-dimensional joint space, we built a 5-nearest-neighbor graph using correlation distance and applied Leiden community detection (resolution = 10) to intentionally oversplit clusters so that small programs could be distinguished. This procedure yielded 187 clusters. We assigned program labels by manual curation, considering each cluster’s mean expression pattern across the global cell-state manifold and gene-set enrichments. During curation, clusters with shared expression patterns and similar perturbation responses—likely oversplits—were merged. Clusters lacking expression patterns matching clearly definable cell types and without significant gene-set overrepresentation were labeled “mixed cell types” and excluded from downstream analyses. Finally, we cleaned the member genes of each gene expression program by computing the mean vector of the member genes in the PCA space and filtering genes by the Pearson correlation coefficient between each gene’s PCA vector and the mean PCA vector of the program. Genes with correlation coefficients less than 0.7 were removed from the programs.

### Compute gene expression program scores

For each program *P* with member genes {*g*_1_, …, *g*_n_p__}, we defined the program score as the mean expression of its members scaled by 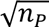:

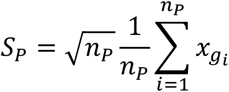

The 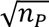 factor down-weights the dependence of the score’s variance on program size, facilitating comparison of score shifts across programs. Under a null model in which gene-level shifts are independent with equal variance, this scaling yields program-score shifts with approximately size-invariant standard deviation.

### Learning functional embeddings of perturbed genes by a mixture-of-experts model

We trained a mixture-of-experts (MoE) model to learn a global functional embedding for perturbed genes from knockdown Perturb-seq datasets. Conceptually, Perturb-seq reports the transcriptomic response to loss of a gene with particular molecular functions. We assume that each gene *g* can be represented by a low-dimensional vector *v_g_* ∈ ℝ_*M*_ in the functional embedding space such that, for a given starting cell state with SCMG embedding *z*, the perturbation-induced shift in readout genes Δ*y* ∈ ℝ_*N*_ is approximately a linear map of *v_g_*. In the MoE formulation, the cell-state-specific linear map is expressed as a convex combination of *K* basis (expert) matrices {*W_k_* ∈ ℝ_*N,M*_}_*k*=1,..,*K*_:

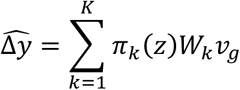

where the gating network takes the SCMG cell-state embedding *z* as input and outputs nonnegative weights *π_k_*(*z*) that sum to one. Intuitively, cells deploy a limited number of response “modes” to perturbations, each represented by an expert, and different cell types mix these modes to varying degrees. This architecture allows the model to capture shared pathway effects across screens while accommodating cell type-specific rewiring of responses. We trained the functional-embedding MoE with embedding dimension *M* = 16, *K* = 8 experts, and *N* = 18108 readout genes (the standard gene set of SCMG). Note that here K is the number of experts, but not the same as the number of shots used for few-shot prediction.

### Clustering functional embeddings of perturbed genes to define perturbation classes

We clustered perturbed genes in the MoE-derived functional space to identify pathway-level modules, which we termed perturbation classes. We included 7 perturb-seq datasets and focused on 608 genes whose perturbations produced reproducible transcriptomic shifts between two K562 perturb-seq dataset (day 6 vs day 8; Pearson *r* > 0.4). In the functional-embedding space, we constructed a 5–nearest-neighbor graph using cosine distance and applied Leiden community detection (resolution = 5), yielding 50 initial clusters. This high resolution aimed to oversplit modules to capture small groups. We then performed manual curation—examining overrepresented gene sets and proximity in the embedding—to merge clusters with similar functional signatures.

### Few-shot prediction of perturbation effects

We performed few-shot prediction using the function embeddings of perturbed gene learned by a mixture-of-experts model (see the “ Learning functional embeddings of perturbed genes by a mixture-of-experts model” section) trained on previously published CRISPRi Perturb-seq datasets (excluding the dataset for prediction evaluation). For example, to evaluate the predict accuracy of the perturbation responses in K562 cells, K562 perturb-seq datasets are withheld from training. For a specified number of shots (K), we selected training perturbations by running K-means clustering on the functional embeddings of all genes perturbed in the evaluation dataset (e.g. K562 screens) and choosing K genes at the cluster centroids to optimize coverage of the embedding space. Using the selected K training perturbations, we fitted a ridge-regression model that takes the functional embeddings of these K perturbed genes as input and optimized the model such that it accurately predicts the perturbation-induced pseudobulk expression-shift vectors of these training genes. We used ridge-regression as the few-shot learning model because it is a simple linear model with regularized weights, suitable for fitting transfer learning tasks with small numbers of data points. We then used this model to predict the perturbation-induced expression-shift vectors of the other genes from their functional embeddings. Performance was assessed as the Pearson correlation between predicted and experimentally pseudobulk gene-expression shift vectors, with the K training perturbations held out from evaluation.

### CRISPRi Perturbation Library Cloning for hESC Perturb-seq Experiment

sgRNA libraries were generated with established methods, with several modifications. Oligonucleotide pools (**Supplementary Table 6**) were ordered from Twist and amplified by PCR using common adapter sequences. The libraries were purified and concentrated with SPRI beads and digested overnight with BstXI and BlpI, alongside an mU6-sgRNA EF1a-GFP sgRNA vector. The digested library was size-selected with an acrylamide gel and the digested vector was size-selected with an agarose gel, and the DNA was extracted. The digested vector and digested library were ligated overnight at 16°C with high-concentration T4 ligase (NEB). The ligation product was purified with a Clean and Concentrate column (Zymo) and transformed into Mega-X cells (ThermoFisher) and grown directly in liquid culture overnight, and then megaprepped. A small sample was taken before liquid culture and plated to confirm coverage; coverage was maintained at >1000 colonies per library element.

### Lentivirus generation and titering

Lentivirus for the Perturb-seq experiments was generated with established methods^110^ and titered empirically by infecting H1 hESC cells with increasing quantities of lentivirus and measuring the fraction of GFP+ cells by flow cytometry three days later.

### Balancing screen

A balancing screen was done to even the representation of sgRNAs at the point of Perturb-seq analysis. We aimed to measure the growth phenotypes for each sgRNA so that we could overrepresent sgRNAs that decrease growth in our Perturb-Seq perturbation libraries. CRISPRi dropout screens with a genome-wide sgRNA library were conducted largely according to established methods. Inducible CRISPRi H1 hESCs were cultured in StemFlex (ThermoFisher) and passaged with Accutase (StemCell) every third day, with ROCKi included in the medium the day after passage. The cells were transduced at >1000x coverage and <30% transduction rate, as assessed by flow cytometry three days later. Three days after transduction, the cells were selected with puromycin for three days (in the presence of ROCKi for the first two days) and then allowed to recover for two days. Zim3 dCas9 was induced by the addition of 25 nM doxycycline and the cells were maintained in culture at >1000x coverage. Samples were collected before induction and every 2-3 days afterwards. We also collected samples from the day-11 time point, fixed them, and stained them with antibodies against Oct4, Nanog, and Sox2, and then used FACS to separate triple-positive cells from cells that were negative in at least one of the three markers. DNA was extracted, amplified, and sequenced according to standard methods^110^.

### Perturb-seq library design

All sgRNAs for the Perturb-seq experiment were chosen from the CRISPRiV2 and CRISPRiV2.1 libraries. We strove to generate an sgRNA library that targeted most genes that gave transcriptional phenotypes when knocked down in hESCs, selected based on the logic that most genes with transcriptional phenotypes also cause growth phenotypes^57^ or impact the expression of pluripotency markers. To this end, we identified sgRNAs with any evidence for growth phenotypes in our balancing screen, sgRNAs with any evidence for altered pluripotency marker expression, and selected all genes with at least two sgRNAs showing evidence for phenotype. In total, we targeted 2978 genes with two sgRNAs each. We also included 370 negative control sgRNAs. Drawing on our balancing screen, we over-represented sgRNAs with strong growth phenotypes in our oligo library, such that sgRNAs expected to drop out 4-fold over the time course of the Perturb-seq experiment were 4-fold over-represented in the initial library, etc.

### Perturb-seq experiment

Inducible CRISPRi H1 hESCs were cultured in StemFlex (ThermoFisher) and passaged with Accutase (StemCell) every third day, with ROCKi included in the medium the day after passage. The cells were transduced at >1000x coverage and <20% transduction rate, as assessed by flow cytometry three days later. The cells were selected by puromycin for three days (in the presence of ROCKi for the first two days) and allowed to recover for two days. Zim3-dCas9 was induced by the addition of 25 nM doxycycline and the cells were maintained in culture at >1000x coverage (accounting for sgRNA over-representation in the initial library). Eight days after Zim3-dCas9 induction, the cells were passaged with accutase, sorted for GFP-positive signal, and processed by 5’ scRNA-seq (10X Genomics) with spike-in primer against the sgRNA constant region to enable 5’ direct capture^111^.

### Processing and sgRNA calling of the hESC perturb-seq data

We used the cellranger 8.0.1 to process the sequenced reads, which counted the UMIs of endogenous mRNAs and the barcodes of sgRNAs. We filtered the cells to only keep cells with at least 500 genes with non-zero counts and targeted by only one sgRNA with at least 20 barcode counts.

### Calculation of pseudo-bulk gene-expression shifts

We filtered the perturbed genes for the calculation of pseudo-bulk expression profiles. First, we selected the perturbed genes that were detected in at least 50 cells. Then we ensured that the perturbed genes were significantly down-regulated by requiring that the mean expression level of the perturbed gene was less than 90% of the mean level of cells with non-targeting guides and the p-value of one-sided Mann–Whitney U tests was less than 0.01. For each selected perturbed gene, we calculated the pseudo-bulk gene-expression profile as the mean expression profile across all cells receiving the sgRNA targeting the perturbed gene. We subset the readout genes to the standard gene set defined in the “Define a standard gene set for human and mouse” section. Then we normalized the sum of each pseudo-bulk expression profile to 10,000 and log1p transformed the profile. For each perturbed gene, we defined the perturbation-induced gene-expression shift vector as the difference between the normalized pseudo-bulk expression profile of cells receiving the sgRNA targeting the gene and the normalized profile of the cells receiving non-targeting control sgRNAs. We noticed that a small fraction of perturbed genes yielded large transcriptional shifts, but the shift vectors from the two guide RNAs targeting the same gene do not correlate well. We filtered out the perturbations with the norm of gene expression shift vector greater than 4 and the correlation between the shift vectors for the two guide RNAs less than 0.2. This filtering removed 259 genes.

### Gene-set over-representation analysis

We performed gene-set over-representation analysis to help annotate the gene-expression programs, perturbation classes, as well as the single-cell clusters of the newly generated hESC perturb-seq dataset. We used the GSEApy package^112^ to perform gene-set over-representation analysis and gene-set enrichment analysis. We chose the gene sets defined in the GO_Biological_Process_2023, Reactome_2022, CORUM and KEGG_2021_Human libraries^65–68^ to perform the analyses.

We used the enrichr function of GSEApy to perform gene-set over-representation analysis. The enrichr function takes a list of genes as input and computes a p-value for each gene set in the selected gene-set libraries, indicating whether the gene set is significantly overrepresented in the input list of genes.

### Single-cell clustering of the hESC perturb-seq dataset

We performed two rounds of single-cell clustering of the hESC perturb-seq dataset. In the first round, we normalized the total gene count for each cell to 10,000 and performed log1p transformation. Then we subset the genes to be the genes with strong expression shifts identified by pseudo-bulk analysis. Specifically, we computed the mean expression shift of each readout gene induced by each perturbed gene (averaged across all cells receiving the sgRNA targeting the same perturbed gene) and selected the readout genes that had absolute expression shift values being greater than 0.2 for at least two perturbed genes to generate the gene expression profile of each cell. Then we z-score-scaled the gene expressions, performed dimensionality reduction by PCA, calculated a 20-nearest-neighbor graph, and clustered the cells by the Leiden algorithm. For each first-round cluster, we visualized the cells by UMAP and if the UMAP showed that the cluster could be further separated into distinct groups, we performed a second-round clustering with the same method.

### Identification of perturbations enriched in single-cell clusters or classes

We calculated the perturbations enriched in single-cell groups at both the cluster level and the class level, where the class is a group of clusters, such as the germ-layer differentiation class. For a given granularity level, we calculated a few metrics to evaluate the enrichment of each sgRNA in each cell group. For each cell group, we counted the number of cells carrying the sgRNA of interest in that group. We calculated the foreground-vs-background fraction ratio as the fold change in the fractions of cells with the sgRNA of interest in the cell group over the fraction of cells with the sgRNA of interest outside the cell group. We calculated the p-value of the fold change with the chi-squared test. We adjusted the p-values for multiple hypothesis correction by the Benjamini–Hochberg procedure. We defined that a sgRNA was enriched in a cell group if there were at least two cells with the sgRNA in that group; the foreground-vs-background fraction ratio was greater than 2; and the adjusted p-value was less than 0.001. We defined that a perturbed gene was enriched in a cell group if it had at least one sgRNA enriched in the group and another sgRNA with at least two cells in the group and the foreground-vs-background ratio being greater than 1. To make the dot plot of perturbed genes enriched in cell groups, we treated different sgRNAs targeting the same gene as the same perturbation and computed the enrichment ratios and p-values with the same method as described above.

## REFERENCES

1. Hrovatin, K. et al. Considerations for building and using integrated single-cell atlases. Nat. Methods 22, 41–57 (2025).

2. Rood, J. E. et al. The Human Cell Atlas from a cell census to a unified foundation model. Nature 637, 1065–1071 (2025).

3. Rozenblatt-Rosen, O. et al. The Human Tumor Atlas Network: Charting tumor transitions across space and time at single-cell resolution. Cell 181, 236–249 (2020).

4. Ecker, J. R. et al. The BRAIN Initiative Cell Census Consortium: Lessons learned toward generating a comprehensive brain cell atlas. Neuron 96, 542–557 (2017).

5. Jain, S. et al. Advances and prospects for the Human BioMolecular Atlas Program (HuBMAP). Nat. Cell Biol. 25, 1089–1100 (2023).

6. Hawrylycz, M. et al. A guide to the BRAIN Initiative Cell Census Network data ecosystem. PLoS Biol. 21, e3002133 (2023).

7. Bunne, C. et al. How to build the virtual cell with artificial intelligence: Priorities and opportunities. Cell 187, 7045–7063 (2024).

8. Baek, S., Song, K. & Lee, I. Single-cell foundation models: bringing artificial intelligence into cell biology. Exp. Mol. Med. 57, 2169–2181 (2025).

9. Ahlmann-Eltze, C., Huber, W. & Anders, S. Deep-learning-based gene perturbation effect prediction does not yet outperform simple linear baselines. Nat. Methods 22, 1657–1661 (2025).

10. Sun, S., Zhu, J., Ma, Y. & Zhou, X. Accuracy, robustness and scalability of dimensionality reduction methods for single-cell RNA-seq analysis. Genome Biol. 20, 269 (2019).

11. Luecken, M. D. et al. Benchmarking atlas-level data integration in single-cell genomics. Nat. Methods 19, 41–50 (2022).

12. Xiao, Y. et al. Gene Perturbation Atlas (GPA): a single-gene perturbation repository for characterizing functional mechanisms of coding and non-coding genes. Sci. Rep. 5, 10889 (2015).

13. Qiu, C. et al. A single-cell time-lapse of mouse prenatal development from gastrula to birth. Nature 626, 1084–1093 (2024).

14. Qiu, C. et al. Systematic reconstruction of cellular trajectories across mouse embryogenesis. Nature Genetics vol. 54 328–341 Preprint at 10.1038/s41588-022-01018-x (2022).

15. Suo, C. et al. Mapping the developing human immune system across organs. Science 376, eabo0510 (2022).

16. Cao, J. et al. A human cell atlas of fetal gene expression. Science 370, eaba7721 (2020).

17. Sikkema, L. et al. An integrated cell atlas of the lung in health and disease. Nat. Med. 29, 1563–1577 (2023).

18. Reed, A. D. et al. A single-cell atlas enables mapping of homeostatic cellular shifts in the adult human breast. Nat. Genet. 56, 652–662 (2024).

19. Yao, Z. et al. A high-resolution transcriptomic and spatial atlas of cell types in the whole mouse brain. Nature 624, 317–332 (2023).

20. Tabula Sapiens Consortium* et al. The Tabula Sapiens: A multiple-organ, single-cell transcriptomic atlas of humans. Science 376, eabl4896 (2022).

21. Han, X. et al. Construction of a human cell landscape at single-cell level. Nature 581, 303–309 (2020).

22. Litviňuková, M. et al. Cells of the adult human heart. Nature 588, 466–472 (2020).

23. Elmentaite, R. et al. Cells of the human intestinal tract mapped across space and time. Nature 597, 250–255 (2021).

24. Bhaduri, A. et al. An atlas of cortical arealization identifies dynamic molecular signatures. Nature 598, 200–204 (2021).

25. Domínguez Conde, C., et al. Cross-tissue immune cell analysis reveals tissue-specific features in humans. Science 376, eabl5197 (2022).

26. Park, J.-E. et al. A cell atlas of human thymic development defines T cell repertoire formation. Science 367, eaay3224 (2020).

27. Lake, B. B. et al. An atlas of healthy and injured cell states and niches in the human kidney. Nature 619, 585–594 (2023).

28. Arutyunyan, A. et al. Spatial multiomics map of trophoblast development in early pregnancy. Nature 616, 143–151 (2023).

29. La Manno, G. et al. Molecular architecture of the developing mouse brain. Nature 596, 92–96 (2021).

30. Tabula Muris Consortium. A single-cell transcriptomic atlas characterizes ageing tissues in the mouse. Nature 583, 590–595 (2020).

31. Hrovatin, K. et al. Delineating mouse β-cell identity during lifetime and in diabetes with a single cell atlas. Nat. Metab. 5, 1615–1637 (2023).

32. Eraslan, G. et al. Single-nucleus cross-tissue molecular reference maps toward understanding disease gene function. Science 376, eabl4290 (2022).

33. Kuppe, C. et al. Spatial multi-omic map of human myocardial infarction. Nature 608, 766–777 (2022).

34. Yu, Q. et al. Charting human development using a multi-endodermal organ atlas and organoid models. Cell 184, 3281–3298.e22 (2021).

35. Jardine, L. et al. Blood and immune development in human fetal bone marrow and Down syndrome. Nature 598, 327–331 (2021).

36. He, P. et al. A human fetal lung cell atlas uncovers proximal-distal gradients of differentiation and key regulators of epithelial fates. Cell 185, 4841–4860.e25 (2022).

37. Fawkner-Corbett, D. et al. Spatiotemporal analysis of human intestinal development at single-cell resolution. Cell 184, 810–826.e23 (2021).

38. Efthymiou, V. et al. Single-nucleus analysis of human white adipose tissue reveals adipocyte subsets with distinct metabolic profiles. bioRxivorg 2025.09.14.673351 (2025) doi:10.1101/2025.09.14.673351.

39. Allen, W. E., Blosser, T. R., Sullivan, Z. A., Dulac, C. & Zhuang, X. Molecular and spatial signatures of mouse brain aging at single-cell resolution. Cell 186, 194–208.e18 (2023).

40. Lengyel, E. et al. A molecular atlas of the human postmenopausal fallopian tube and ovary from single-cell RNA and ATAC sequencing. Cell Rep. 41, 111838 (2022).

41. Vento-Tormo, R. et al. Single-cell reconstruction of the early maternal-fetal interface in humans. Nature 563, 347–353 (2018).

42. Cowan, C. S. et al. Cell types of the human retina and its organoids at single-cell resolution. Cell 182, 1623–1640.e34 (2020).

43. Wiedemann, J. et al. Differential cell composition and split epidermal differentiation in human palm, sole, and hip skin. Cell Rep. 42, 111994 (2023).

44. Enge, M. et al. Single-cell analysis of human pancreas reveals transcriptional signatures of aging and somatic mutation patterns. Cell 171, 321–330.e14 (2017).

45. Tyser, R. C. V. et al. Single-cell transcriptomic characterization of a gastrulating human embryo. Nature 600, 285–289 (2021).

46. Yanagida, A. et al. Naive stem cell blastocyst model captures human embryo lineage segregation. Cell Stem Cell 28, 1016–1022.e4 (2021).

47. McInnes, L., Healy, J. & Melville, J. UMAP: Uniform Manifold Approximation and Projection for Dimension Reduction. arXiv [stat.ML] (2018).

48. Trimm, E. & Red-Horse, K. Vascular endothelial cell development and diversity. Nat. Rev. Cardiol. 20, 197–210 (2023).

49. Lopez, R., Regier, J., Cole, M. B., Jordan, M. I. & Yosef, N. Deep generative modeling for single-cell transcriptomics. Nat. Methods 15, 1053–1058 (2018).

50. Theodoris, C. V. et al. Transfer learning enables predictions in network biology. Nature 1–9 (2023).

51. Cui, H. et al. scGPT: toward building a foundation model for single-cell multi-omics using generative AI. Nat. Methods 21, 1470–1480 (2024).

52. Heimberg, G. et al. A cell atlas foundation model for scalable search of similar human cells. Nature 638, 1085–1094 (2025).

53. Rosen, Y., et al. Universal Cell Embeddings: A Foundation Model for Cell Biology. bioRxiv 2023.11.28.568918 (2023) doi:10.1101/2023.11.28.568918.

54. Burclaff, J. et al. A proximal-to-distal survey of healthy adult human small intestine and colon epithelium by single-cell transcriptomics. Cell. Mol. Gastroenterol. Hepatol. 13, 1554–1589 (2022).

55. Travaglini, K. J. et al. A molecular cell atlas of the human lung from single-cell RNA sequencing. Nature 587, 619–625 (2020).

56. Treutlein, B. et al. Dissecting direct reprogramming from fibroblast to neuron using single-cell RNA-seq. Nature 534, 391–395 (2016).

57. Replogle, J. M. et al. Mapping information-rich genotype-phenotype landscapes with genome-scale Perturb-seq. Cell 185, 2559–2575.e28 (2022).

58. Adamson, B. et al. A multiplexed single-cell CRISPR screening platform enables systematic dissection of the unfolded protein response. Cell 167, 1867–1882.e21 (2016).

59. Frangieh, C. J. et al. Multimodal pooled Perturb-CITE-seq screens in patient models define mechanisms of cancer immune evasion. Nat. Genet. 53, 332–341 (2021).

60. Tian, R. et al. Genome-wide CRISPRi/a screens in human neurons link lysosomal failure to ferroptosis. Nat. Neurosci. 24, 1020–1034 (2021).

61. Tian, R. et al. CRISPR interference-based platform for multimodal genetic screens in human iPSC-derived neurons. Neuron 104, 239–255.e12 (2019).

62. Jiang, L. et al. Systematic reconstruction of molecular pathway signatures using scalable single-cell perturbation screens. Nat. Cell Biol. 27, 505–517 (2025).

63. Nakatake, Y. et al. Generation and Profiling of 2,135 Human ESC Lines for the Systematic Analyses of Cell States Perturbed by Inducing Single Transcription Factors. Cell Rep. 31, 107655 (2020).

64. Traag, V. A., Waltman, L. & van Eck, N. J. From Louvain to Leiden: guaranteeing well-connected communities. Scientific Reports vol. 9 Preprint at 10.1038/s41598-019-41695-z (2019).

65. Aleksander, S. A. et al. The Gene Ontology knowledgebase in 2023. Genetics 224, (2023).

66. Gillespie, M. et al. The reactome pathway knowledgebase 2022. Nucleic Acids Res. 50, D687–D692 (2022).

67. Giurgiu, M. et al. CORUM: the comprehensive resource of mammalian protein complexes-2019. Nucleic Acids Res. 47, D559–D563 (2019).

68. Kanehisa, M., Furumichi, M., Sato, Y., Matsuura, Y. & Ishiguro-Watanabe, M. KEGG: biological systems database as a model of the real world. Nucleic Acids Res. 53, D672–D677 (2025).

69. Pfrieger, F. W. & Ungerer, N. Cholesterol metabolism in neurons and astrocytes. Prog. Lipid Res. 50, 357–371 (2011).

70. Genaro-Mattos, T. C., Anderson, A., Allen, L. B., Korade, Z. & Mirnics, K. Cholesterol biosynthesis and uptake in developing neurons. ACS Chem. Neurosci. 10, 3671–3681 (2019).

71. Duan, Y. et al. Regulation of cholesterol homeostasis in health and diseases: from mechanisms to targeted therapeutics. Signal Transduct. Target. Ther. 7, 265 (2022).

72. Hubert, K. A. & Wellik, D. M. Hox genes in development and beyond. Development 150, dev192476 (2023).

73. Brooks, E. R. & Wallingford, J. B. Multiciliated cells. Curr. Biol. 24, R973–82 (2014).

74. McNab, F., Mayer-Barber, K., Sher, A., Wack, A. & O’Garra, A. Type I interferons in infectious disease. Nat. Rev. Immunol. 15, 87–103 (2015).

75. Sedger, L. M. & McDermott, M. F. TNF and TNF-receptors: From mediators of cell death and inflammation to therapeutic giants - past, present and future. Cytokine Growth Factor Rev. 25, 453–472 (2014).

76. Jacobs, R. A., Jordan, M. I., Nowlan, S. J. & Hinton, G. E. Adaptive mixtures of local experts. Neural Comput. 3, 79–87 (1991).

77. Do, B. T. et al. Nucleotide depletion promotes cell fate transitions by inducing DNA replication stress. Dev. Cell 59, 2203–2221.e15 (2024).

78. Muhr, J. & Hagey, D. W. The cell cycle and differentiation as integrated processes: Cyclins and CDKs reciprocally regulate Sox and Notch to balance stem cell maintenance. Bioessays 43, e2000285 (2021).

79. Pakos-Zebrucka, K. et al. The integrated stress response. EMBO Rep. 17, 1374–1395 (2016).

80. Kroemer, G., Mariño, G. & Levine, B. Autophagy and the integrated stress response. Mol. Cell 40, 280–293 (2010).

81. Pflaum, J., Schlosser, S. & Müller, M. P53 family and cellular stress responses in cancer. Front. Oncol. 4, 285 (2014).

82. Metcalf, M. G., Higuchi-Sanabria, R., Garcia, G., Tsui, C. K. & Dillin, A. Beyond the cell factory: Homeostatic regulation of and by the UPRER. Sci. Adv. 6, eabb9614 (2020).

83. Law, J. C., Ritke, M. K., Yalowich, J. C., Leder, G. H. & Ferrell, R. E. Mutational inactivation of the p53 gene in the human erythroid leukemic K562 cell line. Leuk. Res. 17, 1045–1050 (1993).

84. Read, A. & Schröder, M. The unfolded protein response: An overview. Biology (Basel) 10, 384 (2021).

85. An, J. et al. Identification of spliceosome components pivotal to breast cancer survival. RNA Biol. 18, 833–842 (2021).

86. Lempiäinen, H. & Shore, D. Growth control and ribosome biogenesis. Curr. Opin. Cell Biol. 21, 855–863 (2009).

87. Petibon, C., Malik Ghulam, M., Catala, M. & Abou Elela, S. Regulation of ribosomal protein genes: An ordered anarchy. Wiley Interdiscip. Rev. RNA 12, e1632 (2021).

88. Destefanis, F., Manara, V. & Bellosta, P. Myc as a regulator of ribosome biogenesis and cell competition: A link to cancer. Int. J. Mol. Sci. 21, 4037 (2020).

89. van Riggelen, J., Yetil, A. & Felsher, D. W. MYC as a regulator of ribosome biogenesis and protein synthesis. Nat. Rev. Cancer 10, 301–309 (2010).

90. George, S., Heng, B. C., Vinoth, K. J., Kishen, A. & Cao, T. Comparison of the response of human embryonic stem cells and their differentiated progenies to oxidative stress. Photomed. Laser Surg. 27, 669–674 (2009).

91. Scaramuzzino, L. et al. Uncovering the metabolic and stress responses of human embryonic stem cells to FTH1 gene silencing. Cells 10, 2431 (2021).

92. Mandal, S., Lindgren, A. G., Srivastava, A. S., Clark, A. T. & Banerjee, U. Mitochondrial function controls proliferation and early differentiation potential of embryonic stem cells. Stem Cells 29, 486–495 (2011).

93. Loh, K. M., Lim, B. & Ang, L. T. Ex uno plures: molecular designs for embryonic pluripotency. Physiol. Rev. 95, 245–295 (2015).

94. Adomavicius, T. et al. The structural basis of translational control by eIF2 phosphorylation. Nat. Commun. 10, 2136 (2019).

95. Diao, S. et al. Lineage plasticity of the integrated stress response is a hallmark of cancer evolution. bioRxivorg 2025.02.10.637516 (2025) doi:10.1101/2025.02.10.637516.

96. Marcucci, F. & Rumio, C. How tumor cells choose between epithelial-mesenchymal transition and autophagy to resist stress-therapeutic implications. Front. Pharmacol. 9, 714 (2018).

97. Norman, T. M. et al. Exploring genetic interaction manifolds constructed from rich single-cell phenotypes. Science 365, 786–793 (2019).

98. Srivatsan, S. R. et al. Massively multiplex chemical transcriptomics at single-cell resolution. Science 367, 45–51 (2020).

99. Alda-Catalinas, C. et al. A single-cell transcriptomics CRISPR-activation screen identifies epigenetic regulators of the zygotic genome activation program. Cell Syst. 11, 25–41.e9 (2020).

100. DeMeo, B. et al. Active learning framework leveraging transcriptomics identifies modulators of disease phenotypes. Science 390, eadi8577 (2025).

101. Huang, K. et al. Sequential optimal experimental design of perturbation screens guided by multi-modal priors. bioRxiv 2023.12.12.571389 (2023) doi:10.1101/2023.12.12.571389.

102. Stuart, T. & Satija, R. Integrative single-cell analysis. Nature Reviews Genetics vol. 20 257–272 Preprint at 10.1038/s41576-019-0093-7 (2019).

103. Triana, S. et al. Single-cell proteo-genomic reference maps of the hematopoietic system enable the purification and massive profiling of precisely defined cell states. Nat. Immunol. 22, 1577–1589 (2021).

104. Dyer, S. C. et al. Ensembl 2025. Nucleic Acids Res. 53, D948–D957 (2025).

105. Wolf, F. A., Alexander Wolf, F., Angerer, P. & Theis, F. J. SCANPY: large-scale single-cell gene expression data analysis. Genome Biology vol. 19 Preprint at 10.1186/s13059-017-1382-0 (2018).

106. Bruford, E. A. et al. Guidelines for human gene nomenclature. Nat. Genet. 52, 754–758 (2020).

107. Loshchilov, I. & Hutter, F. Decoupled weight decay regularization. arXiv [cs.LG] (2017).

108. Abdulla, S. et al. CZ CELLxGENE Discover: a single-cell data platform for scalable exploration, analysis and modeling of aggregated data. Nucleic Acids Res. 53, D886–D900 (2024).

109. Peidli, S. et al. scPerturb: harmonized single-cell perturbation data. Nat. Methods 21, 531–540 (2024).

110. Gilbert, L. A. et al. Genome-scale CRISPR-mediated control of gene repression and activation. Cell 159, 647–661 (2014).

111. Replogle, J. M. et al. Combinatorial single-cell CRISPR screens by direct guide RNA capture and targeted sequencing. Nat. Biotechnol. 38, 954–961 (2020).

112. Fang, Z., Liu, X. & Peltz, G. GSEApy: a comprehensive package for performing gene set enrichment analysis in Python. Bioinformatics 39, btac757 (2023).

